# Sex-determining 3D regulatory hubs revealed by genome spatial auto-correlation analysis

**DOI:** 10.1101/2022.11.18.516861

**Authors:** Irene Mota-Gómez, Juan Antonio Rodríguez, Shannon Dupont, Oscar Lao, Johanna Jedamzick, Ralf Kuhn, Scott Lacadie, Sara Alexandra García-Moreno, Alicia Hurtado, Rafael D. Acemel, Blanche Capel, Marc A. Marti-Renom, Darío G. Lupiáñez

**Affiliations:** Max-Delbrück Center for Molecular Medicine in the Helmholtz Association (MDC), Berlin Institute for Medical Systems Biology (BIMSB), Epigenetics and Sex Development Group, Berlin, Germany; CNAG-CRG, Centre for Genomic Regulation (CRG), Barcelona Institute of Science and Technology (BIST), Baldiri i Reixac 4, 08028 Barcelona, Spain; Department of Cell Biology, Duke University Medical Center, Durham, NC, USA; Institut de Biologia Evolutiva (UPF-CSIC), Department of Medicine and Life Sciences, Universitat Pompeu Fabra, Parc de Recerca Biomèdica de Barcelona, Barcelona, Spain; Max-Delbrück Center for Molecular Medicine in the Helmholtz Association (MDC), Berlin, Germany; Max-Delbrück Center for Molecular Medicine in the Helmholtz Association (MDC), Berlin Institute for Medical Systems Biology (BIMSB), Computational Regulatory Genomics Group, Berlin, Germany; Centre for Genomic Regulation (CRG), Barcelona Institute of Science and Technology (BIST), Dr. Aiguader 88, 08003 Barcelona, Spain; Universitat Pompeu Fabra (UPF), 08002 Barcelona, Spain; ICREA, 08010 Barcelona, Spain

## Abstract

Mammalian sex is determined by opposing networks of ovarian and testicular genes that are well characterized. However, its epigenetic regulation is still largely unknown, thus limiting our understanding of a fundamental process for species propagation. Here we explore the 3D chromatin landscape of sex determination *in vivo*, by profiling FACS-sorted embryonic mouse gonadal populations, prior and after sex determination, in both sexes. We integrate Hi-C with ChIP-seq experiments using *METALoci*, a novel genome spatial auto-correlation analysis that identifies 3D enhancer hubs across the genome. We uncover a prominent rewiring of chromatin interactions during sex determination, affecting the enhancer hubs of hundreds of genes that display temporal- and sex-specific expression. Moreover, the identification of the 3D enhancer hubs allows the reconstruction of regulatory networks, revealing key transcription factors involved in sex determination. By combining predictive approaches and validations in transgenic mice we identify a novel *Fgf9* regulatory hub, deletion of which results in male-to-female sex reversal with the upregulation of ovarian-specific markers and the initiation of meiosis. Thus, spatial auto-correlation analysis is an effective strategy to identify regulatory networks associated to biological processes and to further characterize the functional role of the 3D genome.

## INTRODUCTION

Reproduction is a fundamental aspect of life that depends on the differentiation of compatible sexes. In mammals, sex is determined by a complex but tightly balanced network of ovarian- and testicular-promoting factors^1^. Prior to sex determination, gonads from both sexes are bipotential, as they can either develop as ovaries or testes. In XY individuals, the *Sex-determining region Y protein* (*Sry)* gene is sufficient to tilt this balance: its expression in the supporting lineage of the bipotential gonad results in the activation of its direct downstream target, the pro-testicular gene *SRY-box transcription factor 9* (*Sox9*). Subsequently, SOX9 interacts with the *fibroblast growth factor 9* (FGF9) morphogen to propagate the male-determining signal to the entire gonad, thus suppressing ovarian-specific genes and promoting testicular development. In the absence of *Sry* expression, ovarian development takes place, with the activation of several members of the *Wnt* pathway, such as *R-spondin 1* (*Rspo1*), *Wnt family member 4* (*Wnt4*) and *catenin beta 1* (*Ctnnb1*). Subsequently, sex-determining signals induce dramatic changes on cell differentiation, hormone synthesis and, ultimately, a physical and behavioral transformation of the entire organism. While decades of research have revealed the identity of multiple genes associated to sex determination, the epigenetic regulatory aspects of this process are still unclear.

In vertebrates, gene expression is controlled by the action of cis-regulatory elements (CREs), which serve as binding platforms for transcription factors (TF)^2^. CREs provide spatio-temporal specificity to transcription, acting in cooperation to constitute complex and pleiotropic expression patterns during development. However, to exert their function, CREs may enter into physical proximity with their target genes, a process mediated by the 3D folding of the chromatin. The development of chromosome conformation capture methods, in particular Hi-C^3^, revealed that vertebrate genomes fold into distinct levels of organization^4–6^. At the megabase scale, genomes segregate into active (A) and inactive (B) compartments, which reflect the clustering of loci according to their transcriptional state. At the submegabase scale, genomes organize into topologically associating domains (TADs), which represent large genomic regions with increased interaction frequencies containing genes and their putative CREs. Non-coding mutations, either affecting CRE function or TAD organization, can result in developmental diseases or cancer^7–9^. Moreover, they can serve as a driving force for species adaptation^10–12^.

Non-coding mutations have been also associated to variations in sex determination. For example, large duplications and deletions in the surrounding genomic desert of the *Sox9* gene, and including the eSR-A enhancer, have been identified in human patients affected with sex reversal^13^. Importantly, XY mice carrying a deletion of the homologous genomic region (Enh13) develop ovaries instead of testes^14^. More recently, a large inversion has been associated with ovotesticular development in female moles^10^. This structural variant incorporates active enhancers into the TAD of the *Fgf9* gene which, in contrast to other mammals, is upregulated during early female gonadogenesis in moles. These experimental evidences delineate a critical role for epigenetic regulation in sex determination. While recent studies have provided an initial exploration of these important aspects^15,16^, the lack of information on three-dimensional (3D) chromatin organization has limited progress beyond the molecular dissection of selected loci. This has a direct impact in our capacity to genetically diagnose Differences of Sex Development (DSD), a group of conditions that alter reproductive capacities in humans^17^.

Here, we explore the 3D regulatory landscape of mammalian sex determination *in vivo*. By combining FACS-sorting and low-input Hi-C^18^, we generate high-resolution chromatin interaction maps of the mouse gonadal supporting lineage, before and after sex determination, in both sexes. Additionally, we introduce *METALoci*, a novel computational approach that integrates Hi-C and epigenetic data to provide an unbiased quantification of the regulatory environment around each gene in the genome. Using *METALoci* in a comparative approach, we further reveal prominent 3D environment changes in hundreds of genes that display sex- and temporal specificity. We subsequently reconstruct regulatory networks associated to sex determination, which include key known factors as well as novel regulators. We further employ *METALoci* as a predictive tool to identify regulatory regions across the non-coding genome without prior knowledge. We validate these predictions by investigating the regulation of the pro-testicular gene *Fgf9*, identifying a critical region that leads to male-to-female sex reversal upon deletion in transgenic mouse models. Our method highlights the important role of 3D genome organization in guiding sex determination, a process of critical relevance for species reproduction.

## RESULTS

### Changes in compartments and TADs are moderate during sex determination

We explored the 3D regulatory landscape of sex determination in the gonadal supporting lineage of both sexes, prior to and after its commitment to the female or male fate (that is, four states named XX E10.5, XX E13.5, XY E10.5, and XY E13.5, **Fig. 1a**). For this purpose, we employed mouse lines expressing cell-specific markers that allow the isolation of gonadal populations by fluorescence-activated cell sorting (FACS). Progenitor supporting cells were isolated from both sexes at E10.5, using a *Sf1-eGFP* line^19^. At this stage, prior to the expression of *Sry*, these cells are bipotential due to their capacity to differentiate towards either the female or male lineage. At E13.5, a *Sox9-eGFP* line was used to isolate Sertoli cells from developing testes^19^, while a *Runx1-eGFP* line was employed to obtain their counterparts in ovaries, the granulosa cells^20^.

**Fig. 1:**
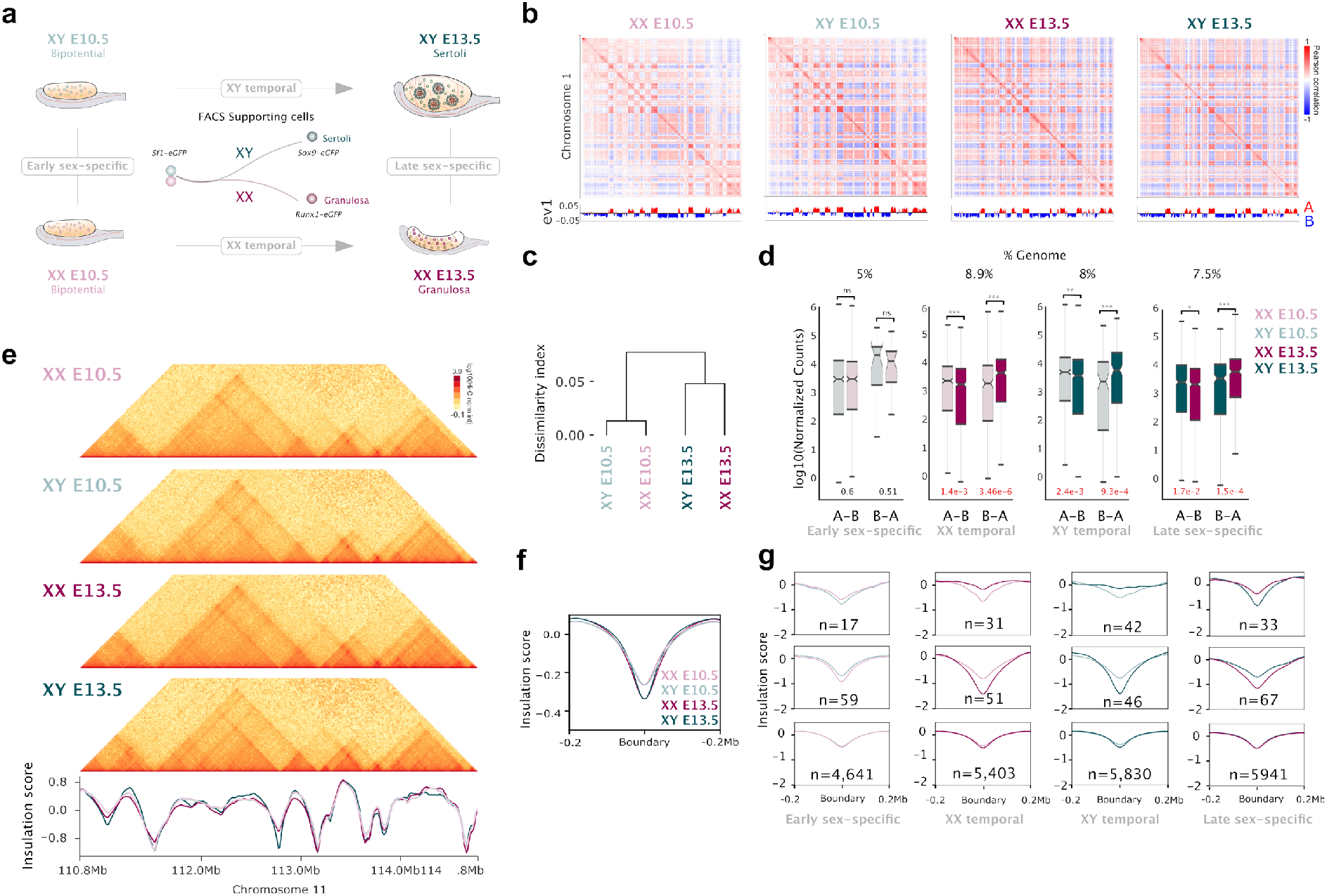
Moderate changes in compartment and TAD organization during sex determination. ***a**. Experimental setup of FACS-sorted gonadal populations. **b**. Compartment analyses showing eigenvectors for chromosome 1 in different samples. Eigenvectors are computed from the matrices of the physical interaction between pairs of loci across the chromosome. **c**. Dissimilarity index in A/B compartments between different samples. **d**. Gene expression levels for genes that switched compartments. Note that changes in gene expression occur in all comparisons, except in XY E10.5 vs. XX E10.5. **e**. Upper panel. Hi-C maps of the Sox9 locus in different samples. Lower panel. Insulation scores for the same genomic region **f**. Insulation score at TAD boundaries. Note the increase in insulation during the transition from bipotential to Sertoli or granulosa cells. **g**. Pairwise comparison of insulation scores at boundaries. The two upper rows correspond to boundaries that change insulation between samples, while the lower row depicts stable boundaries. Note that stable boundaries are more abundant in any comparison.*

3D chromatin interactions from these isolated cell populations were profiled at high resolution, using a low-input Hi-C protocol^18^ and generating between 750 and 950 million valid pairs per sample (**Extended Data Table 1**). We subsequently aimed to identify potential changes in 3D chromatin organization during sex determination by employing standard Hi-C analyses. First, we identified compartments at 100Kb resolution, which resulted in a high correlation between biological replicates (**Fig. 1b** and **Extended Data Fig. 1**). As expected, A compartments were enriched in H3K27ac, open chromatin and increased gene density, in contrast to B compartments (**Extended Data Fig. 2**). Across samples, the proportion assigned to A compartments fluctuated between 42-50% and 50-58% for B compartments (**Extended Data Fig. 3**). Yet, dissimilarity index analyses revealed an increased compartment correlation between male and female cells prior to sex determination (**Fig. 1c; Extended Data Fig. 4**), thus reflecting higher similarities at the bipotential stage. These correlations decreased after sex is determined and cells progress towards the granulosa or Sertoli cell fate. We next identified switches in compartments by performing pairwise comparisons between samples (**Extended Data Fig. 5**). Overall, we observed that compartment switches involved 7.4-8.9% of the genome and correlated well with expected changes in gene expression (*i.e*., A to B: decreased expression; B to A: increased expression; **Fig. 1d**; **Extended Data Fig. 5**). An exception was observed in the comparison between the female and male bipotential stage, in which only 5.3% of the genome varied (**Extended Data Fig. 5**). Since *Sry* is still not active at this developmental timepoint, the sexual dimorphism in compartments may be induced by the different X chromosome complement between the sexes^21^. Interestingly, this early sex-specific variation in compartments was not associated to changes at the transcriptional level (**Fig. 1d**). This may suggest that variations in 3D chromatin organization might precede changes in gene expression, as previously described for other differentiation processes^22^.

Next, we focused on exploring changes at the level of TAD organization. For each sample, we calculated insulation scores and identified a total of 6,179 TAD boundaries (**Fig. 1e**). Metaplot analysis revealed that insulation at boundary regions increased during the transition from bipotential to the differentiated stages (**Fig. 1f**), as also described in other biological systems^23^. Pairwise comparisons revealed that only 1.49-1.84% of TAD boundaries changed their insulation significantly between sexes or timepoints (**Fig. 1g**). However, manual inspection revealed that most of these changes resulted from quantitative changes in insulation rather than *de novo* formation or disappearance of TAD boundaries (**Extended Data Fig. 6**). In summary, our analyses revealed a moderate variation in 3D chromatin organization during sex determination, reflected by a high degree of conservation in TAD structures and compartment changes that increased as differentiation progressed.

### METALoci *reveals a prominent rewiring of 3D enhancer hubs in a temporal and sex-specific fashion*

Our analyses of TAD dynamics suggests that, at a large scale, 3D chromatin organization is preformed prior to sex determination and maintained afterwards. However, the variation in compartments suggests that certain genomic regions acquire a different epigenetic status during this process, which could indicate changes in regulation occurring at other scales. However, these types of changes are difficult to detect with conventional strategies for Hi-C data analysis, as they rely on the accurate identification of genome structures such as compartments, TADs or loops (*i.e*., highly interacting loci). Therefore, these approaches may not detect other chromatin organization changes that could be of relevance for biological processes such as gene regulation. To overcome these limitations, we developed *METALoci*, an unbiased approach that allows a quantification of regulatory activity loci-by-loci without prior assumptions from the data.

*METALoci* relies on spatial autocorrelation analysis, classically employed in geostatistics^24,25^ to describe how the variation of a variable depends on space at a global and local scales (*e.g*., identifying contamination hotspots within a city^26^). We specifically repurposed this type of analysis to quantify gene regulation, based on the fact that CREs and their target genes may cluster together within the 3D nuclear space and display similar epigenetic properties. Briefly, the overall flowchart of *METALoci* consists of four steps (**Fig. 2a**). First, a genome-wide Hi-C normalized matrix is taken as input and the top interactions selected (**Fig. 2b**).Second, the selected interactions are used to build a graph layout (equivalent to a physical map) using the Kamada-Kawai algorithm^27^ with nodes representing bins in the Hi-C matrix and the 2D distance between the nodes being inversely proportional to their normalized Hi-C interaction frequency (**Fig. 2b**). Third, epigenetic/genomic signals, measured as coverage per genomic bin (*e.g*., ChIP-seq signal for H3K27ac), are next mapped into the nodes of the graph layout (**Fig. 2c**). The fourth and final step involves the use of a measure of autocorrelation (specifically, the Local Moran’s I or LMI^24,25^) to identify nodes and their neighborhoods (that is, other genomic bins within a specified 2D distance in the graph layout) with similar epigenetic/genomic signals (**Fig. 2c**). *METALoci* categorizes each genomic bin according to its signal status as well as that of its surrounding neighborhood (**Fig. 2d**). Specifically, a genomic bin categorized as High-High (HH) will be enriched for the signal, but also other bins that are in spatial proximity with it (**Fig. 2d**). In contrast, bins marked as Low-Low (LL) represent those that are depleted of the signal in both the corresponding bin and its spatial neighborhood. High-Low (HL) and Low-High (LH), will then represent bins that are enriched in signal, but not their neighborhood, and vice versa. Finally, the group of genomic bins that are spatially contiguous and statistically significant for the spatial enrichment of the signal (*i.e*., those classified by *METALoci* as significant HH bins, including their direct neighbors in the graph layout **Fig. 2d**, red outlined shape) are here named “metaloci”. It is important to note that *METALoci* quantifies the spatial autocorrelation of the input signal for each genomic bin, which facilitates the direct comparison between different datasets.

**Fig. 2:**
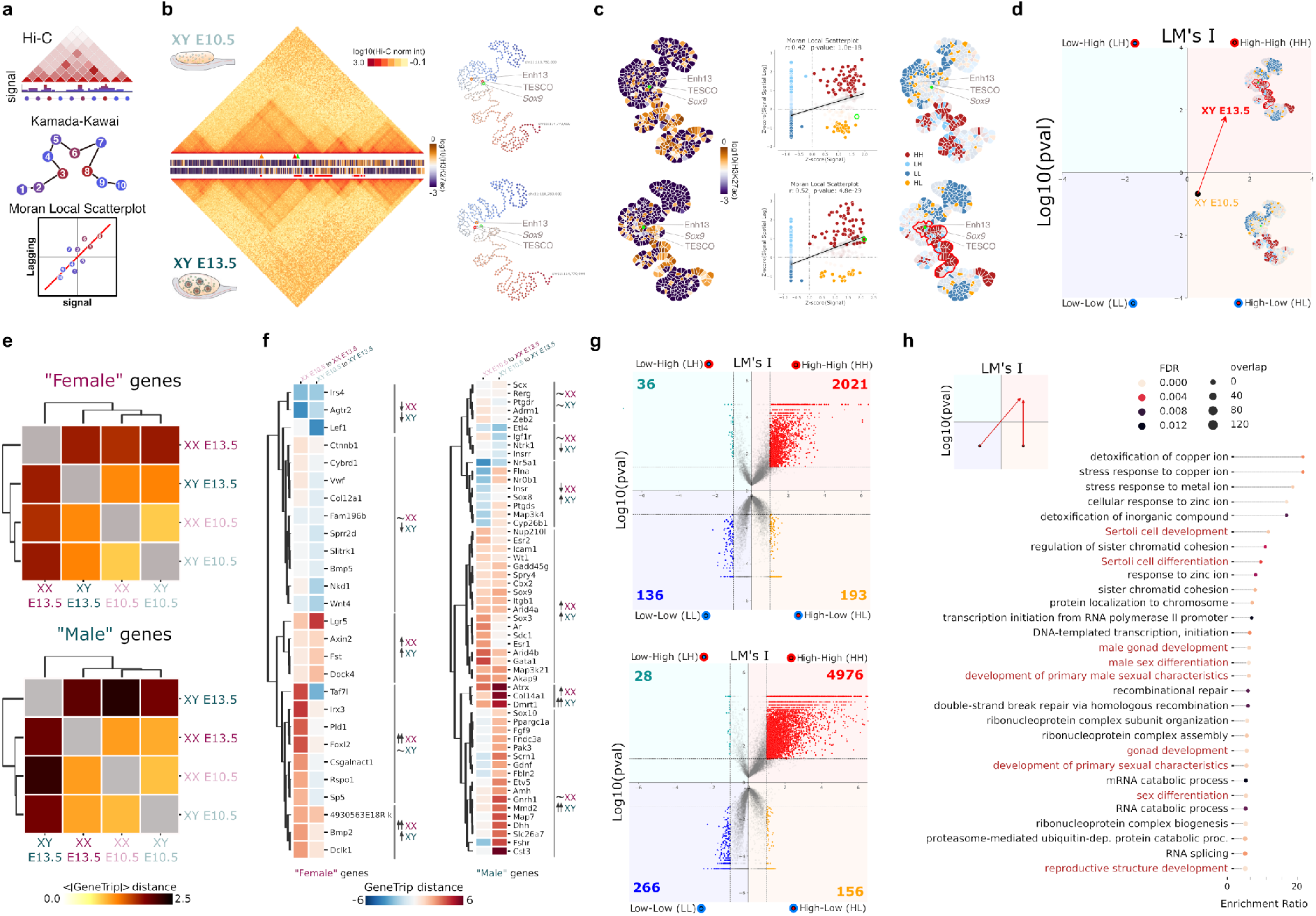
METALoci captures extensive rewiring of 3D enhancer hubs during sex differentiation. ***a**. Schematic METALoci pipeline (detailed in **Methods**) **b***. *Left. Hi-C data and H3K27ac signal for* Sox9 *locus centered at chr11:110,780,000-114,770,000 coordinates. Hi-C and H3K27ac ChIP-seq for XY E10.5 and XY E13.5 cells are displayed. The position of the* Sox9 *promoter (green arrowhead), as well as the Enh13 (orange arrowhead) and TESCO (red arrowhead), are highlighted. Squared red marks under H3K27ac track indicate the non-continuous metaloci detected for* Sox9 *locus at XY E13.5. Right. Taking as input Hi-C data, a 2D layout is generated using the Kamada-Kawai algorithm. The layout highlights the* Sox9 *locus (green circle) as well as the Enh13 (orange circle) and the TESCO (red circle) enhancers. **c**. Left. H3K27ac signal is mapped into the graph layout and represented as a “Gaudí plot” (detailed in **Methods**). Middle. LMI scatter plot where each point representing a node in the graph layout is placed within the 4 quadrants of the LMI (that is, HH, LH, LL, and HL). Points with solid color are statistically significant (p<0.05). The point of the node containing the* Sox9 *locus is highlighted with a green circle. Right. Gaudí plot highlighting in space the classification of each bin into the LMI quadrants with solid color indicating statistical significance (p<0.05)*. Sox9 *HH metaloci outlined in red. **d**. LMI transition (here called “gene trip”) for the* Sox9 *locus from a HL non-significant to a significant HH enhancer hub during the differentiation of Sertoli cells (XY E10.5 to XY E13.5). A gene trip is the length (in arbitrary units) of the vector connecting LM’s I and p-value coordinates between two time points. In the example*, Sox9 *gene trip (red arrow) was 2.29. A positive gene trip indicates that the resulting vector points towards the HH quadrant. A negative gene trip indicates that the vector points towards the LL quadrant. **e**. Mean absolute gene trip for genes acquiring female and male-specific expression during sex differentiation^28^. The gene trips are larger for “male” genes in XY cells upon differentiation compared to “female” genes in XX cells. **f**. Individual gene trips for each female and male-specific genes during sex-determination. Genes are grouped by unsupervised clustering based on gene trips from E10.5 to E13.5 in female and male differentiation. **g**. LMI quadrants for XY E10.5 (top) and XY E13.5 cells (bottom) for 24,027 annotated gene promoters in the mm10 reference genome. Quadrants include the total number of statistically significant genes in each quadrant. Note the increased numbers after differentiation in the HH and LL, denoting simultaneous activation and repression of genes during differentiation. **h**. GO Biological Process enrichment analysis for genes that transition from LL or HL to HH during Sertoli cell differentiation*.

For each gene of the mouse genome, we applied *METALoci* to reconstruct its 3D enhancer hubs (or metaloci) during sex determination. With that purpose, we integrated our Hi-C datasets with H3K27ac ChIP-seq signal^16^, which marks active promoters and enhancers. Thus, HH metaloci for H3K27ac can be considered 3D hubs of enhancers activating the expression of the resident genes. To assess the accuracy of *METALoci* in detecting such enhancer hubs we first analyzed the *Sox9* locus, which is directly activated by SRY and is essential to trigger the male pathway and Sertoli cell differentiation^14^. *METALoci* results recapitulated known changes in the regulation of the *Sox9* gene (**Fig. 2b-d**). At XY E10.5, the *Sox9* promoter remained in an inactive status and was not associated to a HH metaloci. In contrast, at XY E13.5 when Sertoli cells differentiate, the *Sox9* gene changed its regulatory status to HH, with both its promoter and environment characterized as enriched for H3K27ac signal (**Fig. 2c**). Those changes parallel the transcriptional dynamics of *Sox9*, which is expressed at low levels at the bipotential stage in both sexes, but subsequently activated in male Sertoli cells^19^. Importantly, *METALoci* captured the dynamic interaction of the *Sox9* promoter with two known enhancers, TESCO^29^ and Enh13^14^ (**Fig. 2b; Extended Data Fig. 7**), which are essential for its male-specific upregulation, Sertoli cell differentiation and the testicular program. Furthermore, *METALoci* also predicts the existence of additional regulators downstream of the *Sox9* promoter (**Fig. 2c**). Globally, *Sox9* transition (here called “gene trip” and detailed in **Methods**) within the four quadrants of the *METALoci* analysis stated its HL to HH trip between E10.5 and E13.5 in XY cells, in agreement with its known activation (**Fig. 2d**). Conversely, *METALoci* also captured regulatory changes associated with female differentiation. In particular, the promoter of *Bmp2*, which was classified as LL at early stages, gained an active enhancer environment in granulosa cells (HH) being part of a large contiguous patch of H3K27ac enrichment from two spatial proximal metaloci, which included a recently described enhancer element^16^ (**Extended Data Fig. 8**).

To further assess *METALoci* capacity to identify known sex-determination biology, we curated primarily from literature^28^ a list of 27 and 55 genes associated to female and male gonad development, respectively. The analysis of these subsets of genes with dimorphic expression during sex differentiation revealed changes in regulation upon differentiation in both sexes. During the differentiation from bipotential to granulosa cells, most female-specific genes acquired an active enhancer environment, while their male-specific counterparts lost it (**Extended Data Fig. 9**). The opposite trend was observed during the differentiation of Sertoli cells, with the acquisition of active regulatory activity at most male-specific genes and a loss in their female-specific counterparts (**Extended Data Fig. 9**). Further analysis revealed that the magnitude of the calculated gene trips, measured as mean absolute length of the vectors connecting the gene state at E10.5 and E13.5 (**Extended Data Fig. 9**), in sex-specific genes was moderate at early stages, but increased upon differentiation (**Fig. 2e**). Interestingly, the changes in regulatory environment were more prominent for male-specific than for female-specific genes during the differentiation into the corresponding sex. A more detailed comparison of each individual gene trip revealed distinct mechanisms governing how sex-specific genes acquire their dimorphic expression pattern (**Fig. 2f**). Most male genes (46 out of 55) gain an active regulatory environment during Sertoli cell differentiation. In contrast, this mechanism was not as common for female-specific genes during granulosa cell differentiation (14 out of 27). Interestingly, the loss in regulatory activity for genes upregulated in the opposite sex was higher during male differentiation (13 out of 27), than during female differentiation (8 out of 55). Overall, these results suggest that the male differentiation program is sustained by more pronounced changes in regulation than the female program.

We next explored genome-wide changes in 3D enhancer hubs during sex determination. We observed that, independently of the sex, the number of genes categorized as HH doubled as differentiation progressed, thus reflecting the activation of specific transcriptional programs (**Fig. 2g**). Concomitantly, genes categorized as LL also doubled, suggesting the repression of additional pathways. Functional enrichment analyses revealed that genes transitioning from HL or LL towards HH during Sertoli cell differentiation (XY E10.5 to XY E13.5) were associated with relevant terms for this biological process (**Fig. 2h**). Specifically, we observed an over-representation (False Discovery Rate, FDR <0.01) of biological terms related to Sertoli cell differentiation, male sex development and RNA processing. While RNA processing signatures were also present, an enrichment in sex-specific processes was not observed during female differentiation (XX E10.5 to XX E13.5) (**Extended Data Fig. 10**), which could reflect the molecular similarity of granulosa cells with their bipotential progenitors at this developmental stage^19^.

Overall, *METALoci* revealed a prominent rewiring of 3D enhancer hubs during sex determination, affecting the regulatory landscape of hundreds of genes (**Extended Data File 1**). Importantly, such changes could not be systematically captured with conventional Hi-C analyses, thus highlighting the sensitivity of *METALoci* in capturing changes that are meaningful for gene regulation.

### Reconstruction of gene regulatory networks uncovers novel genes involved in sex determination

Next, we sought to identify which regulatory networks of transcription factors (TF) were associated to the 3D enhancer hubs discovered by *METALoci*. To reconstruct these networks, we intersected ATAC-seq peaks from these FACS-sorted cell types^16^ with the identified H3K27ac HH metaloci. This integration allowed us to narrow down the genomic location of CREs for those genes likely to be actively transcribed within the broader regions defined by *METALoci*. Next, we performed transcription factor footprinting analyses on the ATAC-seq peaks within metaloci using TOBIAS^30^, which allowed the association of CREs with their predicted bound TFs. This analysis revealed distinct gene regulatory networks for each sample (**Fig. 3a**), in which known sex-determining TFs display a high degree of connectivity (that is, a given TF sequence motif is found several times within the accessible peaks of the target TF metaloci, and vice versa). For example, WT1 or NR5A1 (SF1) appeared as ubiquitously represented in both sexes during all stages, consistent with their essential role in the early formation of the bipotential gonad^31,32^. Other sex-determining TF increased their degree of connectivity in the networks as differentiation progressed, such as SOX9, SOX10 and DMRT1 in Sertoli, or RUNX1 and GATA4 in granulosa cells (**Fig. 3a**).

**Fig. 3.**
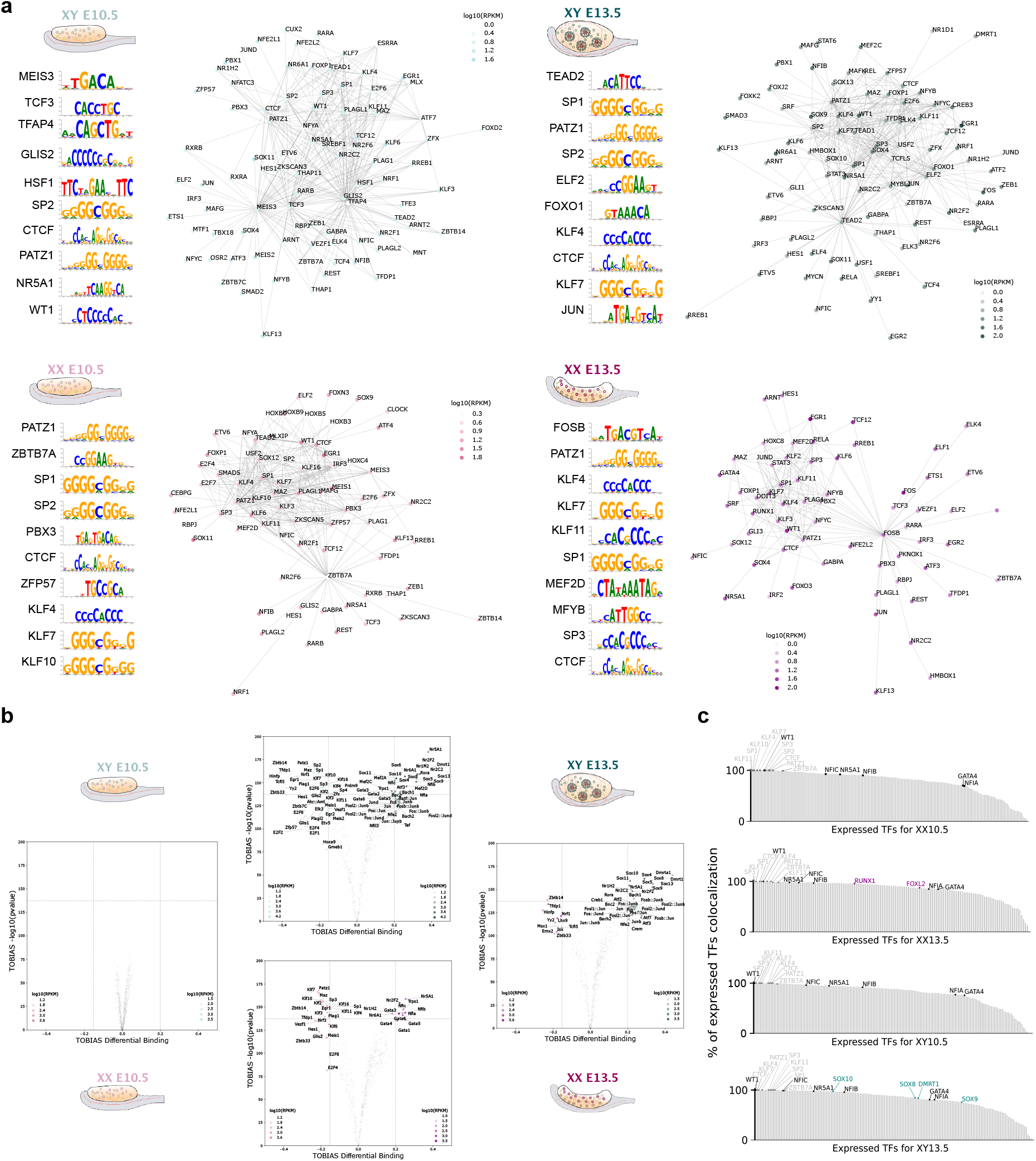
Gene regulatory networks uncover novel genes involved in sex determination. ***a**. Gene regulatory networks for the four analyzed samples displaying TFs with more than 10 connections. The top 10 TFs with the highest number of connections at their corresponding binding motif matrices are displayed at the left of each plot. **b**. TF differential binding analysis performed in pairwise comparisons using TOBIAS^30^. Note the absence of significantly differentially-bound TF between XY10.5 and XX10.5. **c**. STRIPE analysis for colocalized TFs. For each cell type, the bar graph shows all expressed TFs in the cell type sorted by the percentage of TFs that colocalize with them. A TF is considered to colocalize if both are found together in the genome at least 20 instances. TF names are colored according to being universal (grey), gonad-specific (black), female specific (purple), and male specific (dark green).*

To further explore the hierarchy of sex-determining TFs, we selected the top 10 factors with the highest degree of connectivity within each network, highlighting a total of unique 26 TF for the four analyzed samples. As expected, CTCF was ubiquitously enriched in all samples (**Fig. 3a**), consistent with its crucial role in 3D chromatin organization^33^. Furthermore, this analysis revealed a common enrichment in GC-rich motifs, such as PATZ1 or members of the Specificity-Protein/Krüppel-like factor (SP/KLF) family. Interestingly, this group of TFs render chromatin accessible to other TFs^34^. In addition, other TF motifs resulted in increased connectivity in specific samples, such as PBX3 in female bipotential cells. While this factor has not been previously associated to sex determination, data from the international mouse phenotyping consortium (IMPC^35^; www.mousephenotype.org) indicates that *Pbx3* knockout mice develop a female-specific phenotype, characterized by the complete absence of ovaries. Similarly, knockout mice for *Meis3*, which our analysis found to be highly connected in XY bipotential cells, display abnormal testicular development. Other TFs resulted in increased connectivity at later stages, such as FOXO1 specifically enriched in Sertoli cells. The increased connectivity for this TF in the male network contrasts with the known role of Forkhead-box factors in triggering female development^20^. However, although *Foxo1* knockout mice display early lethality, *in vitro* knockdown studies have revealed a role as an upstream regulator of *Sox9* in chondrocyte development^36^. Thus, *METALoci* is able to provide novel insights into genes that compose the sex determination network.

To gain further understanding of the regulatory dynamics of sex determination, we performed differential TF binding analyses at a genome-wide scale. Importantly, this analysis did not identify any significantly differentially-bound TF at the bipotential stage, thus reflecting the molecular similarities between sexes at this developmental timepoint^16,19,37,38^. However, differences in TF binding motifs were prominently observed after sex was determined (**Fig. 3b**). Specifically, the binding for SP/KLF, MEIS or GLIS factors was selectively reduced, while the chromatin binding for GATA, NFI factors or SF1 increased as differentiation progressed, irrespectively of the sex. Furthermore, Sertoli cells acquired sex-specific signatures such as increased binding for SOX factors, DMRT1, as well as the Activator Protein 1 (AP-1, also known as FOS:JUN). Interestingly, AP-1 has been previously shown to mediate 3D regulatory hubs in differentiation processes^39,40^, which is consistent with the increased transcriptional response at the onset of Sertoli cell differentiation, in comparison to granulosa cells^19,37,38^. We further explored the combinatorial nature of TF interaction during sex determination by examining the colocalization of binding motifs within ATAC-seq footprints^34^. The analysis revealed that several TF act as “stripe factors”, which may increase chromatin accessibility for other co-binding TF partners^34^ (**Fig 3c; Extended Data Fig. 11**). While some of these factors have been described as universal (*e.g*., KLF/SP), others like WT1 are specific for gonadal tissue^41^. Altogether, these findings demonstrate that *METALoci* is able to capture regulatory changes that are functionally relevant for sex determination.

### METALoci simulations reveal a novel non-coding region downstream of the Fgf9 gene associated with male-to-female sex reversal

Using *METALoci* and TF foot-printing analysis, we defined the potential location of active gonadal CREs associated to each gene, in a temporal and sex-specific fashion (**Extended Data File 2**). To further validate our findings, we investigated the regulation of *Fgf9*, a pro-testicular gene that encodes a morphogen that upregulates *Sox9* in developing testis and inhibits the female pathway. Consequently, *Fgf9* ablation results in male-to-female sex reversal in transgenic mice^42^. Furthermore, gain in *Fgf9* copy numbers have been identified in patients with 46 XX sex reversal^43^. Yet, despite the critical role of *Fgf9* in controlling sex determination, nothing is known about its regulation.

To identify the critical regions for *Fgf9* gonadal regulation, we computationally scanned the entire locus and estimated the effect of deletions of 50 Kb fragments (that is, 5 contiguous bins of 10 Kb) in the enhancer 3D hubs analyzed by *METALoci* (**Fig. 4a** and **Methods**). In other words, we assessed whether deleting 5 bins at a time in a moving window of 1 bin steps over the studied region would disrupt the *Fgf9* HH metaloci in XY E13.5. Briefly, to do so, we computationally removed the selected bins, their Hi-C interactions, and their H3K27ac signal and re-computed the metaloci for *Fgf9*. If the resulting metaloci was perturbed by the simulated deletion, we considered that the bins removed are important for the HH metaloci of *Fgf9*. Indeed, in XY bipotential cells, when *Fgf9* is expressed at low levels, we observed that critical regulatory regions were predicted to be located proximal to the coding region of the gene. In contrast, in XY Sertoli cells, we observed a switch towards distal regulation that was concomitant with the upregulation of *Fgf9*^19^. Specifically, we identified a non-coding region located approximately 250 Kb downstream of *Fgf9*, whose deletion was predicted to be disruptive of its HH metaloci. Interestingly, the human homologous region contains GWAS hits associated with abnormal testosterone levels^44–46^, a phenotype that is consistent with alterations in *FGF9* expression (**Extended Data Table 2**).

**Fig. 4:**
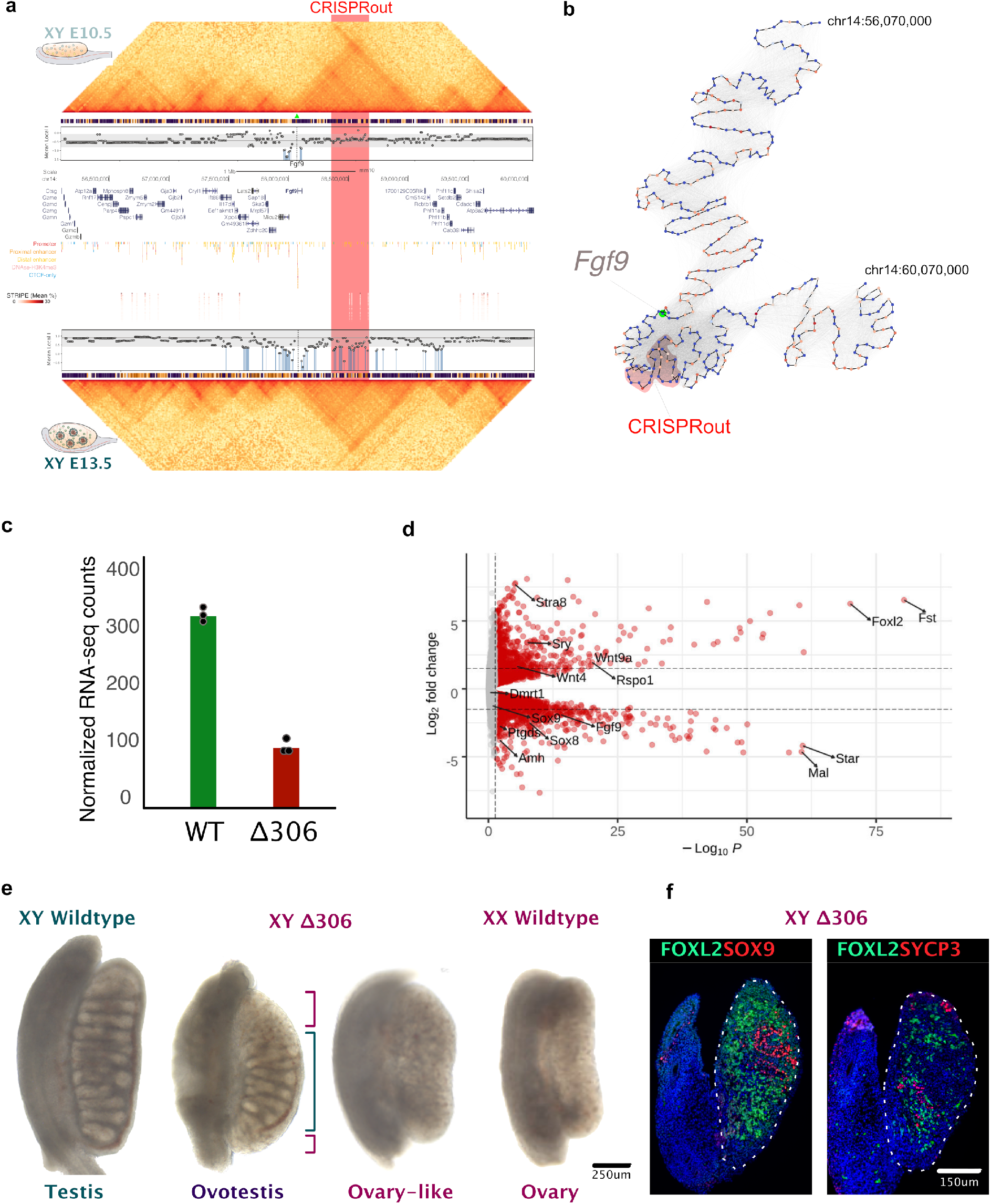
A novel non-coding region at the *Fgf9* locus is associated with male-to-female sex reversal. ***a**. Predictive scanning analysis at the* Fgf9 *locus. Hi-C and ChIP-seq tracks are displayed for XY E10.5 (upper panel) and XY E13.5 (lower panel). Vertical blue lines mark regions whose deletion is predicted to decrease the LM’s I of the metaloci. **b**. Kamada-Kawai layout of the* Fgf9 *locus in XY E13.5. Green dot indicates the bin containing the* Fgf9 *promoter. Red transparent shape indicates deleted region (Δ306). **c**. Expression levels of* Fgf9 *in control wild-type (WT) and mutant (Δ306) gonads. **d**. Volcano plot of RNA-seq from XY E13.5 Δ306 mutant and control gonads. Note downregulation of pro-testicular genes, with the exception of Sry that is upregulated. Also, upregulation of pro-ovarian and meiotic genes. **e**. E14.5 gonads of Δ306 mutants and controls. Note the two phenotypes: ovotestis and ovary-like. Green dashed bracket indicates testicular portion in the center of the ovotestis. Purple bracket indicates ovarian portion at the poles of the ovotestis. **f**. Immunofluorescence on XY E14.5 Δ306 mutants with ovary-like phenotype. Gonads are delineated by a discontinuous line. The ovarian marker FOXL2 (green) marks the presence of granulosa cells. Left. Testicular marker SOX9 indicates the presence of Sertoli cells within testis cords in the center of the gonad. Right. Meiotic marker SYCP3 indicates the initiation of meiosis at the gonadal poles. Note that the gonads corresponding to the ovary-like phenotype in Δ306 mutants may include the presence of a reduced number of testicular cords, as previously described in^42^.*

To validate the *METALoci* predictions, we generated a 306 Kb homozygous deletion (*Δ306*) within the 1.15 Mb TAD of *Fgf9*, in mouse embryonic stem cells (mESC). This deletion included most of the *Fgf9* predicted downstream regulatory region with no annotated genes or regulatory elements in gonads (**Fig. 4a** and **b**). These mESC were subsequently employed to generate transgenic mice via tetraploid complementation assays^47,48^. RNA-seq analyses of XY E13.5 gonads revealed a 2-fold downregulation of *Fgf9* in male mutants compared to controls, associated with the downregulation of other male-specific markers (**Fig. 4c**). Concomitantly, the ovarian program was activated, as reflected by the upregulation of female-specific and the downregulation of male-specific genes (**Fig. 4d**). Meiosis, which is considered to be the first molecular signature of ovarian development^49^, was activated by the upregulation of markers like *Stra8*. Interestingly, *Sry* levels were increased in *Δ306* mutants, which may reflect a transcriptional compensation resulting from impaired testicular development. At E14.5 gonads from XY mutant mice displayed two distinct phenotypes: they either developed as ovotestes or ovary-like gonads (**Fig. 4e**). The ovotestis phenotype was characterized by the development of testicular tissue at the center of the gonad, and ovarian tissue at the poles (**Fig. 4e**). Immunofluorescence analyses confirmed the presence of male markers such as SOX9 in the testicular tissue, as well as female markers like FOXL2 in the ovarian regions of the ovotestis (**Fig. 4f**). The same pattern was observed in ovary-like mutant gonads, although with an increased content in ovarian tissue. These results denoted the initial activation of the male program, but a failure in the propagation of the testis-determining signal to the entire gonad (**Extended Data Fig. 12**). Furthermore, the expression of SYCP3 in mutant gonads confirmed the initiation of the meiotic program observed in RNA-seq experiments (**Fig. 4e** and **f**). Importantly, the two phenotypes observed in *Δ306* mutants mirror those described for the full *Fgf9* knockout^42^, although with differences in their frequency. While an ovarian like phenotype is more often observed in *Fgf9* KO mice, the majority of *Δ306* mutants developed ovotestes, which is consistent with the residual expression of *Fgf9* (**Fig. 4c**). In summary, our transgenic experiments validated the predictive value of *METALoci*, by identifying a novel non-coding region controlling mammalian sex determination.

## DISCUSSION

Many of the current tools for Hi-C comparative analysis rely on predetermined genome structural features such as compartments, TADs or loops. As such, these methods potentially miss other structural features that might be relevant for gene regulation. This is particularly evident for the current study, in which we investigate how the 3D regulatory landscape of sex determination is rewired, transitioning from an initially bipotential system to either one of two alternative fates. Using conventional Hi-C analyses, we observe limited variation in 3D chromatin organization, especially at the TAD level, either between bipotential or differentiated stages. Although this may suggest the existence of a preformed TAD topology, as described for other biological systems^50^, it is in stark contrast with previous studies that demonstrate extensive changes at the transcriptional and epigenetic level during sex determination^16,19,37,38^. Such discrepancy prompted us to develop *METALoci*, an unbiased approach to measure and quantify 3D regulatory activity from Hi-C maps in combination with H3K27ac marks. With this novel approach, we reveal that 3D conformational changes are instead pervasive during sex determination and affect the regulation of hundreds of genes. These changes are minor at the bipotential stage, but are exacerbated as sex is specified and differentiation progresses. Previous transcriptomics analyses have shown that the early supporting lineage of the gonad is primed towards the female fate^19,37,38^, an observation that is consistent with classical theories that postulate this sex as the “default” state^51–53^. Importantly, this female priming is also reflected at a regulatory level, as denoted by the moderate mean *METALoci* gene trips as well as the limited differences in the TF binding landscape during granulosa differentiation, as compared to Sertoli cells. Our analyses also show that regulatory mechanisms that lead to dimorphic gene expression are diverse and locus-specific. During Sertoli differentiation, male-specific genes are commonly associated with an increase in enhancer activity that is concomitant with a decreased activity at many female-specific genes. In contrast, these mechanisms are not as prominently observed during granulosa differentiation. These observations agree with previous studies suggesting that testis differentiation is a more active process than ovarian differentiation, and that this process is also dependent on the repression on the chromatin landscape surrounding female genes in XY gonads^54^. Thus, *METALoci* is particularly suited to capture subtle but meaningful regulatory changes that may be overlooked in conventional Hi-C analysis, providing a new dimension to this type of approach.

In contrast to other biological systems, our understanding of the epigenetic regulation of sex determination is still limited. Here we specifically tackle this issue by reconstructing temporal and sex-specific regulatory networks associated to this process. Our results suggest a three-step model for mammalian sex determination. First, TFs with high affinity for GC-rich sequences, such as members of the SP/KLF family, may remodel chromatin and create an accessible environment for other co-binding partners^34^. As these factors recognize very similar motifs, they might act cooperatively to increase chromatin accessibility through assisted loading^55^. The fact that the binding motifs for these TFs are enriched at the bipotential stage, but reduced after sex is determined, supports their early function. Second, gonad-specific TFs like WT1, SF1 and GATA or NFI factors are recruited to support gonadal outgrowth in both sexes^31,32,56^. Despite the important role of these factors during early gonadal development, our results suggest a fundamental difference in their mechanism of action, with WT1 acting as a gonad-specific stripe factor that renders the chromatin accessible. This tissue-restricted function contrasts with the widespread effects described for other universal stripe factors, like those belonging to the SP/KLF family^34^. Nevertheless, the specific enrichment of all these gonad-specific TFs at E13.5 also highlights additional functions in the subsequent activation of both the female and male-specific programs^57–59^. Finally, sex-determining TFs, such as SOX9 or DMRT1, are recruited to the network to unbalance the bipotential status of the gonad and activate sexual differentiation.

Our limited understanding on how sex determination is regulated represents a major challenge for DSD, for which providing a proper molecular diagnosis is often not possible^60^. In that respect, most of our knowledge on the sex determination process is derived from the identification of DSD-associated mutations from human data. While this approach has been successful in revealing novel candidate genes, it is certainly biased towards the identification of mutations that are compatible with life. Thus, approaches that investigate the regulatory potential of factors may represent an alternative strategy to fill the missing pieces of the sex determination process. Using these approaches, we propose a role in sex determination for SP/KPL and NFI factors during the early steps of gonadal formation. We also uncover a novel role for genes that display a sex bias in regulation like *Pbx3* or *Meis1*, whose disruption has been shown to induce gonadal phenotypes. Interestingly, most of these novel factors are pleiotropic and critical during development, as their inactivation leads to lethality prior to the reproductive stage. Such early lethality might have precluded their previous identification as sex-determining factors using traditional Mendelian disease aproaches and denotes the value of genomic approaches for candidate gene discovery.

A significant percentage of DSD also are expected to result from non-coding mutations that affect gene regulation^60^. Regarding this, *METALoci* allowed us to associate each gene with its temporal and sex-specific 3D regulatory hub, thus providing a functional annotation of the non-coding genome during sex determination. Indeed, our analyses captured regulatory interactions with validated enhancers at the *Bmp2*^16^ and *Sox9* loci^14,29^. In fact, the search for gonadal *Sox9* enhancers has been largely focused on the gene desert that is located upstream of the gene. However, *METALoci* predicts regulatory activity also in the downstream region, which contains enhancers for other tissues like the midbrain^61^ or the jaw^62^. We further demonstrate the accuracy of these predictions, by validating a novel non-coding region at the *Fgf9* locus that controls sex determination in mouse. The gonadal phenotypes from XY mutant mice range from ovotestes to ovaries, thus mimicking those found in human patients carrying coding mutations on *FGF9*^43^, or in *FGFR2* that encodes its gonadal receptor^63^. Remarkably, this novel regulatory region also contains several human GWAS hits associated with fluctuations in testosterone levels^44–46^, a phenotype that is consistent with altered *FGF9* regulation. Overall, *METALoci* revealed important insights into the process of sex determination, going from fundamental mechanisms of gene regulation to their relevance *in vivo*. This highlights the power of integrative genomic approaches to uncover the molecular underpinnings of developmental processes.

## MATERIAL AND METHODS

### Transgenic mice

The *Sf1-eGFP* (*Nr5a1-eGFP*) and *Sox9-eCFP* reporter mouse lines previously generated were maintained on a C57BL/6 (B6) background^64,65^. The *Runx1-GFP* reporter mouse was generously gifted by Dr. Humphrey Yao at NIEHS^20^ (availability at MMRRC_010771-UCD). Timed matings were generated with reporter males and wild-type CD1 females. The morning of a vaginal plug was considered E0.5. Embryos were collected at E10.5 and E13.5, with genetic sex determined using PCR for the presence or absence of the Y-linked gene *Uty* (pF1: TCATGTCCATCAGGTGATGG, pF2: CAATGTGGACCATGACATTG, pR: ATGGACACAGACATTGATGG; two bands indicates XY, one band indicates XX).

### Cell preparation for Hi-C

Gonads were dissected from E10.5 or E13.5 embryos, and the mesonephros was removed using syringe tips. The gonads were incubated in 500ul 0.05% trypsin for 6-10 min at 37°C, then mechanically disrupted in 1X PBS/10% FCS. The cell suspension was pipetted through a 40μm filter top and the supporting cells were collected with fluorescence-activated cell sorting (FACS). After FACS, cells were prepared for Hi-C analysis as follows: cells were spun down at 30×100rpm for 5 minutes at 4°C and resuspended in 250ul 1X PBS/10% FCS. The cells were then fixed in a final concentration of 2% PFA in PBS/10% FCS for 10 minutes at room temperature. The cross-linking reaction was quenched with the addition of 50ul of 1.425M glycine, and the cells were put on ice. Next, the cells were spun 8 minutes, 30×100rpm at 4°C, and the supernatant was removed. The cell pellet was resuspended in cold lysis buffer (50mM TRIS, 150 mM NaCl, 5mM EDTA, 0.5% NP-40, 1.15 TX-100, 6.25X Protease inhibitor cocktail). Cells were centrifuged 3min, 50×1000rpm at 4°C, and the supernatant was removed. Cells were then snap frozen in liquid N2 and stored at −80C until use.

### Hi-C library preparation

Low-input Hi-C protocol was performed from fixed, lysed and snapped frozen cells as previously described^18^, with little modifications. Pelleted aliquots were thawed on ice and resuspended in 25μl 0.5% SDS to permeabilize nuclei and incubated at 62°C for 10’. SDS was quenched by adding 12.5 μl 10% Triton-X-100 and 72.5μl H2O and incubated for 45’at 37°C with rotation. Chromatin was then digested by adding MboI (5U/μl) in two installments in NEB2.1 digestion buffer for a total of 90 minutes, adding the second installment after 45 minutes. Digestion was heat inactivated for 20 minutes at 65°C. DNA overhangs were filled with biotin-14-dATP (0.4mM), dTTP, dGTP, dCTP (10mM) and DNA pol I Klenow (5U/μl) and incubated for 90 minutes at 37°C with gentle rotation. Filled-in chromatin was then ligated by adding ligation master mix (60μl 10X R4 DNA ligase buffer, 50μl 10% Triton-X-100, 6μl BSA (20mg/ml), 2.5μl T4 ligase in 2 installments) for a total of 4 hours at RT with gentle rotation, with the second installment after 2h. Ligated chromatin was then spun down for 5 minutes at 2500g at RT and reversed crosslinked by resuspension in 250μl extraction buffer (10mM Tris pH 8.0, 0.5M NaCl, 1% SDS and 20mg/ml Proteinase K) and incubated for 30 minutes at 55°C while shaking (1000rpm). 56μl 5M NaCl was then added and incubated O/N at 65°C while shaking (1000rpm). Chromatin was then purified by phenol:chloroform:Isoamyl (25:24:1), precipitated with ethanol and resuspended in 15μl Tris pH 8.0. DNA was quantified at this step by Qubit, and RNA was digested by adding 1μl RNase and incubated for 15 minutes at 37°C. Next, biotin was removed from unligated fragments by adding 5μl 10X NEB2 buffer, 1.25μl 1mM dNTP mix, 0.25μl 20mg/ml BSA, 25.5μl H2O and 3μl 3U/μl T4 DNA polymerase and incubated for 4 hours at RT. Sample volume was then brought up to 120μl and DNA shearing was performed with a Covaris S220 (2 cycles, each 50 seconds, 10% duty, 4 intensity, 200 cycles/burst). Biotin pulldown was performed by adding an equal volume of Hi-C sample with Dynabeads MyOne Streptavidin T1 beads (Invitrogen, 65602) and incubated by 15 minutes with rotation. Beads were washed two times with Bind and Wash buffer (10mM Tris-HCl pH 7.5, 1mM EDTA, 2M NaCl) and a final wash with 10mM Tris-HCl pH 8.0 was performed. Samples were resuspended in 50μl Tris-HCl pH 8.0. Library prep was performed using the NEBNext Ultra DNA Library Prep kit for Illumina (E7645L). Briefly, end repair of the libraries was performed by adding 6.5μl 10X End repair reaction buffer and 3μl End Prep Enzyme mix and incubated at RT 30’ followed by 65°C 30 minutes. Next, adaptor ligation was performed by adding 15μl Blunt/TA Ligase master mix and 2.5μl NEBNext adaptor for Illumina and 1μl Ligation enhancer. The mixture was incubated at RT for 15 minutes followed by the addition of 3μl USER Enzyme and an incubation of 15 minutes at 37°C. The beads were separated on a magnetic stand and washed two times with 1X Bind and Wash buffer + 0.1% Triton X-100. Sample was transferred to a new tube and a final wash was performed with 10mM Tris pH8.0 before resuspending the beads in 50μl 10mM Tris pH 8.0.

For PCR library amplification, the sample was divided in 4 reactions of 12.5μl to optimize the number of cycles. The PCR was performed using 12.5μl of Library bound to beads, 25μl 2X NEBNext Ultra II Q5 Master mix, 5μl universal primer 10μM, 5μl indexed PCR primer 10μM and 7.5μl nuclease-free H2O and following the PCR program: 1 minute 98°C 1X, 10 seconds 98°C, 65°C 75 seconds Ramping 1.5°C/sec for 12-20 cycles and 65°C 5 minutes 1X. Double size selection was performed with AmpureXP beads. Library quantification was assessed with Qubit dsDNA HS kit and size and quality of the libraries were checked using TapeStation.

### Hi-C data processing

Raw data was processed and filtered with Juicer^66^ using default parameters. For downstream analysis, Knight-Ruiz normalized Hi-C matrices in hic format were converted to FAN-C^67^ format at 10kb and 100kb resolutions including a low-coverage filtering step, to exclude bins with less than 20% relative coverage. Re-normalization of the filtered matrices was performed using the KR normalization method.

### AB Compartment analysis

AB compartment analysis was performed using FAN-C^67^ in each replicate and chromosome individually in the normalized 100kb KR matrices. After a high Pearson correlation between replicates was confirmed, the matrices of both replicates were merged. The first eigenvector was calculated again in the merged matrices and the sign of the eigenvector was corrected if needed depending on the % of GC and amount ATAC-seq signal in each chromosome independently.

Chromosomes X and Y were excluded from this analysis, since they are not comparable between XX and XY samples. For differential compartment analysis, pairwise comparisons were performed using *bedtools*^68^ by counting number of bins that corresponded to A or B compartment in each sample. Genes and gene expression belonging to each compartment type were included in the analysis using the *bedtools* intersect function. To test significance in differential gene expression between compartments, Benjamini–Hochberg-corrected p-values were reported after pairwise Mann–Whitney U and chi-squared tests.

### Insulation analysis

Insulation scores and boundary scores^69^ were calculated in the 10kb KR normalized, merged matrices using FAN-C^67^ (parameters: window size 500kb, impute_missing= TRUE). To consider that a certain boundary was a TAD boundary, a threshold of 0.25 in the boundary score was used based on visual inspection as recommended by the FAN-C developers. Chromosomes X and Y were excluded for the downstream analysis since they are not comparable between XX and XY samples. A total set of boundaries was obtained using *bedtools cat* (parameters: postmerge=False) and *merge* functions^68^ (parameters: −d=2001). Subsequently, *bedtools intersect* function was used to assess which boundaries were present or absent in each sample.

To generate a quantitative analysis on insulation, pairwise set of boundaries were generated between the samples that needed to be compared (Early sex-specific, Late sex-specific, XX temporal and XY temporal). Next, the insulation scores of both datasets were mapped to the common set of boundaries and an absolute difference in insulation score was calculated. A common z-score for all comparisons was finally calculated. Aggregate profile plots of insulation were generated using *deeptools*^70^.

### METALoci, genome spatial autocorrelation analysis

All *METALoci* analysis was performed using in house developed Python 3 code available as a Jupyter notebook (https://github.com/3DGenomes). The code relies on a series of standard libraries such as SciPy, NumPy (1.21.6), Pandas (1.3.5), Matplotlib (3.5.2), seaborn (0.11.2) as well as other specialized libraries such as GeoPandas (https://geopandas.org, 0.10.2), NetworkX (https://networkx.org, 2.6.3), libpysal (https://pysal.org, 4.6.2), ESDA (https://pysal.org/esda/, 2.4.1), and pyBigWig library from deepTools (https://deeptools.readthedocs.io, 0.3.18).

#### Genome parsing

The first step in *METALoci* is to define the set of genomic regions of interest to analyze. This can be a single gene or a series of *ad-hoc* selected regions. Specifically for this work, the mouse reference genome (mm10, December 2011) was parsed taking as a center point for *METALoci* each of the bins containing a transcription starting site for any of the 24,027 annotated genes. Each region of interest was then centered in its gene TSS, and a total of 2Mb of DNA up- and down-stream was included. This resulted in a list of 24,027 regions of interests each of 4Mb of DNA that were run for the *METALoci* analysis (**Extended Data Table 3**).

#### Hi-C interaction data parsing

*METALoci* uses as input normalized Hi-C interactions at 10Kb resolution, produced as described above. Normalized data was first log_10_ and subset to remove any interaction that was below 1.0 score. This cut-off for interaction selection can be defined by the user and balances the consistency of the resulting Kamada-Kawai layout (next section) and the computational burden. Several cut-offs were assayed for the list of genes, and 1.0 resulted in layouts consistent to others produced with different cut-off with a reasonable computational time. The subset matrix was then transformed from interaction frequencies (*i.e*., a “similarity” matrix) to the inverse of the interactions (*i.e*., a “distance” matrix). Finally, the resulting pair-wise distances between any pair of bins in the region of interest was saved as a sparse matrix to input to the Kamada-Kawai graph layout algorithm.

#### Kamada-Kawai layout

Next, the sparse distance matrix obtained from Hi-C was used as source to generate a graph layout that best represents the observed genomic interactions. This was accomplished by using the Kamada-Kawai graph layout^27^, which attempts to position nodes (that is, genomic bins) on a space of 1 by 1 arbitrary units so that the geometric distance between them is as close as possible to the input distance matrix. To note that the size of the arbitrary space has no effect in the final layout apart from changing its scale, which is irrelevant to the next steps of *METALoci*. The *kamada_kawai_layout* function of the NetworkX python library was used with default parameters to generate the final layouts, as well as obtaining the Cartesian 2D coordinates for each of the genomic bins of 10Kb. Next, the closed Voronoi polygons for each of the bins was calculated using the *Voronoi* function of the SciPy spatial library. The bins at the edge of the layout were closed by placing eight dummy nodes closing the entire space occupied by the layout. This ensured that every single genomic bin had a finite polygon. Next, a buffer distance around each bin was placed corresponding to 1.5 times the mean spatial distance between consecutive genomic bins. Finally, the spatial occupancy of each of the genomic bins was calculated as the intersection of their Voronoi polygon and the buffer space around them. This resulted in a “worm-like” 2D representation of each Kamada-Kawai layout that here we named “Gaudí plots” as they resemble the famous broken tile mosaics or “trencadís” by the Catalan architect Antoni Gaudi (**Fig. 2b**).

#### H3K27ac signal mapping into the graph layout

Next, *METALoci* was input the normalized H3K27ac ChIP-seq signal, produced as described above. H3K27ac coverage per each of the 10Kb bins was obtained using the pyBigWig library, which resulted in a read coverage for each of the bins into the Kamada-Kawai layout. Next, the H3K27ac signal was log_10_ and mapped into each of the polygons of the Gaudí plots. The final result is thus a graph layout representing the input Hi-C interactions and the mapped H3K27ac signal onto the space occupied by each genomic bin. This is then used as input to assess the spatial autocorrelation of H3K27ac using the Local Moran Index approach also known as Local Moran’s I analysis^25^.

#### Local Moran Index autocorrelation analysis

Moran’s I is a measure describing the overall dependence of a given signal over nearby locations in space. Moran’s I is computed as the weighted average of the values of autocorrelation at each *i* sampled point^24,25^:

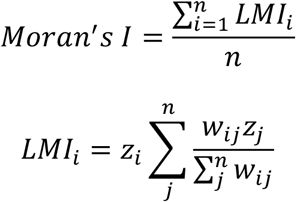

where *z*_i_ is the normalized signal at point *i*, and *w*_ij_ is the assigned weight between point *i* and *j*. Positive LMI are obtained when a point |*z*_i_| > 0 is surrounded by points with similar values (*i.e*., high-high or low-low values), and it is indicative of a hub of points with similar behavior around location *i*. Negative LMI are obtained when |*z*_i_| > 0 and it is surrounded by points with the reverse pattern (*i.e*., high-low or low-high values), and it is suggestive of negative autocorrelation at location *j*. LMI values close to cero indicate poor spatial dependence between contiguous points for the considered signal.

Weights between bins in the Kamada-Kawai graph were calculated based on their spatial distance and the H3K27ac signal. A distance band was assessed based on the *weights.DistanceBand* function of the *libpysal* python library with a distance cut-off corresponding to three times the mean distance between consecutive genomic bins. This ensured that the weights calculated would be based on at least two up- and two down-stream bins as the buffer space for a bin was calculated as 1.5 times the mean distance between consecutive genomic bins (above). Next, the weights were input to the *Moran_Local* function of the ESDA python library with default parameters and for a total of 50,000 permutations to assess the statistical significance of the Moran’s I scores for each bin. The results of the LMI calculations are the Moran’s I score, the Moran’s I quadrant and its significance for each of the bins in the Gaudí plots. Thus, the LMI analysis results in all bins placed into any of the four quadrants of the Moran’s scatter plot. That is, the High-High (HH, red) quadrant for bins that are high in the signal and their neighborhood is also high in signal; the Low-High quadrant (LH, cyan) for bins with low signal but a neighborhood of high signal; the Low-Low quadrant (LL, blue) for bins with low signal in a low-signal neighborhood; and the High-Low signal (HL, orange) for bins that the signal is high but their neighborhood is low. Moreover, after randomizing the signal values over the layout a user-defined times, the algorithm also produces a probability value for each assignment being random. We selected significant HH, LH, LL and HL bins based on a p-value < 0.05. Contiguous bins with significant Moran’s I of the same quadrant and their immediate neighbors correspond to what we call “metaloci” of the signal. Here, we were interested in detecting genes which TSS (*i.e*., the bin in the genomic middle of the layout) was considered a metaloci for enrichment of H3K27ac mark in the HH quadrant (that is, the TSS and its spatial neighborhood are enriched in H3K27ac).

#### Moran’s I volcano plots

Moran’s I inverted volcano graphs (**Fig. 2c**) are plotted by changing the signal of the Moran’s I score for each bin in quadrants LH and LL as well as changing the signal of the Moran’s log_10_(p-value) for bins in quadrants HL and LL. We selected bins containing the TSS gene as significant in each quadrant if the absolute value of the Moran’s I was larger than 1.0 and the absolute log_10_(p-value) larger than 1.3 (p-value < 0.05).

#### Gene trips

A gene trip is calculated as the distance (in arbritary units) that the gene makes in the Moran’s I inverted volcano between two or more sample points. Specifically, here we calculated gene trips for XX and XY cells between time points E10.5 and E13.5. A gene trip is positive if the vector connecting the two analyze time points for the gene of interest points towards the upper-right corner of the Moran’s I inverted volcano (that is, the HH cuadrant). A gene trip is negative if the vector connecting the two analyze time points for the gene of interest points towards the lower-left corner of the Moran’s I inverted volcano (that is, the LL cuadrant).

### GO terms enrichment

Lists of selected genes were used to analyze gene enrichment of Biological Process GO terms using the Web site for WebGestalt (http://webgestalt.org, accessed September 2022)^71^ with coding genes in the mouse genome as a background list. Only GO terms that were deemed significant (False Discovery Rate, FDR < 0.01) were kept.

### Transcription factors analysis

#### TOBIAS transcription factor (TF) binding in each of the analyzed cell types

TOBIAS (https://github.com/loosolab/TOBIAS, 0.13.3)^30^ integrates ATAC-seq footprints with genomic information and (TF) motifs to predict TF binding. We used TOBIAS *ATACorrect* and *FootprintScores* commands with default parameters to correct intrinsic biases of the generated ATAC-seq and to calculate a continuous footprinting score across the genome, respectively. Next, the command *BINDetect* from TOBIAS was used with default parameters to predict specific TF binding by combining the previously generated footprint scores with the information of TF binding motifs from the vertebrates non-redundant JASPAR2022 CORE^72^. Motifs predictions were done exclusively on parts of the genome that were within a metaloci of H3K27ac in each of the four cell types analyzed.

#### TOBIAS TF differential binding between each of the analyzed cell types

The TOBIAS *BINDetect* command with default parameters was also used to compare each one of the analyzed cell types to identify differentially bound transcription factors in metaloci. A TF was considered differentially bound in any of the comparisons if the-log_10_ of TOBIAS p-value was larger than 137.3 and the lower and upper cut-offs for TOBIAS differential binding were smaller than −0.15 and larger than 0.20, respectively. These cut-off values corresponded to the selection of the top 5% differentially binding TF in of the comparisons of the four analyzed cell types.

#### TOBIAS TF networks

TF networks were built by identifying in each of the cell types whether a TF gene metaloci was predicted to be targeted by another TF. For the analysis, we used only TF that were expressed in at least 33% of the single-cells analyzed in previous published dataset^73^ (below). Next, two TF were considered to be linked in the network if one was targeting the gene of the other in at least 10 signatures in its metaloci. Each TF in the network was represented as a node and the edge between them weighted by the number of TF-TF relationships detected by TOBIAS. Next, we used the *kamada_kawai_layout* function of the NetworkX python library to build a network and assess the *degree_counts* function (that is, the number of edges on a node) of NetworkX to assess the top 10 most connected TF in the network. In total, the analysis resulted in 26 unique TF selected in the top most connected for the four cell types analyzed.

#### STRIPE analysis

Using the scRNA-seq profiles at each of the cell types^73^ (below), we selected expressed (RPKM>0) TFs in at least 33% of cells. This resulted in 198, 186, 153, and 176 expressed TFs in XX10.5, XX13.5, XY10.5 and XY13.5, respectively. Next, for each site in the genome corresponding accessible DNA within a metaloci (*i.e*., ATAC-seq peak), we identified shared motifs predicted by TOBIAS for all pairwise comparison of selected TFs. This allowed to identify the so-called “stripe factors” (that is, TFs that share motifs with several other TFs)^34^. Next, the genomic location of these stripe factors was annotated and represented as stripes under the their genomic bin (**Fig. 4a** as example).

### Simulation of genomic perturbations

To computationally predict the effect of CRISPRing out regions of the genome, we devised a strategy where five consecutive bins of 10Kb would be removed using a running window from the beginning to the end of the region of interest in one bin steps. Once a set of 5 bins were removed, all interactions from those bins bin as well as the H3K27ac signal were removed and a new *METALoci* analysis was performed on the resulting Hi-C map and H3K27ac signal. Next, we assessed whether a particular deletion of 50Kb (5 bins) could affect the metaloci status for the bin containing the TSS of the gene of interest. Bin removals that decreased the Moran’s I for the TSS by more than one standard deviation of all analyzed deletions were annotated as predicted perturbation affecting the gene of interest (**Fig. 4**, blue lines in predictions for the *Fgf9* gene).

### Generation of *Fgf9 Δ306* knockout mutants

A deletion of the *METALoci-* predicted region within the *Fgf9* region was generated on G4 mouse embryonic stem cells (mESC), using CRISPR/Cas9 as previously described^48^. Two guide RNAs (sgRNAs) were designed in the regions of interest using Benchling (https://www.benchling.com). The sequences of the sgRNAs were the following: 5’CACCGGCTCCGATAAGATCTGAGC 3’ within the bin number 230 and 5’CACCGTGAGTGCAGCTTTCATCGTA 3’ within the bin number 259 of the *METALoci* perturbation analysis. The absence of the deleted region was assessed by genotyping the flanking regions of the deletion and by genomic qPCR using 3 different pairs of primers located in different areas inside the deletion (**Extended Data Table 4**). Edited cells were then used to generate embryos using tetraploid complementation assay as previously described^47,48^. CD-1 females and males of various ages, were used as donors and fosters for embryo retransferring by tetraploid aggregation. The specimens isolated to perform experimental analysis were Bl6/129Sv5 male, E13.5 and E14.5 in age. All mice were housed in standard cages at the Animal Facility of the Max-Delbrück Center for Molecular Medicine in Berlin in a pathogen-free environment.

### Immunofluorescence

Gonads were dissected out at E14.5, fixed in Serra fixative solution (70% EtOH, 30% FA, 10% Acetic acid), prepared for standard histological methods with paraffin embedding and sectioned in 5μm slides. Immunofluorescence was performed as previously described^10^. The primary antibodies and working dilutions used in this study were: SOX9 (Merck millipore, AB5535, 1:600), FOXL2 (abcam, ab5096-100ug, 1:150) and SCYP3 (abcam, ab15093, 1:200). The secondary antibodies and working dilutions were: Alexa Fluor 488 donkey anti goat IgG (life technologies, A11055, 1:200), Alexa Fluor 555 donkey anti rabbit IgG (life technologies, A31572, 1:200).

### RNA-seq

XY *Fgf9 Δ306* KO and XY control (wild-type, WT) gonads were dissected at the stage of E13.5 in 1xPBS and snap frozen in liquid nitrogen. RNA was then extracted from individual gonads using RNeasy Micro Kit (Qiagen, 74004), following the manufacturer’s specifications. Quality of RNA was assessed using TapeStation and samples were stored a maximum of 1 week at −80C. Libraries were prepared using the NEBNext Ultra II Directional RNA library prep kit for Illumina (E7760), using the protocol that included the poly(A) Magnetic isolation Module (E7490) following the specifications of the manufacturer. Library quality was checked in a Tapestation. Sequencing was performed at 200xPE in a NovaSeq 6000 sequencer.

### ATAC-seq processing

ATAC-seq reads were obtained from GEO (GSE871155)^16^ and trimmed for adapters using *flexbar*^74^ (parameters: -u 10) followed by mapping to the mm10 genome assembly with *bowtie2* with default parameters^75^. Mapped reads were filtered for mapping quality and PCR duplicates using *samtools view* and *markdup*^76^ (parameters: -q 30). The resulting BAM files were converted to BED files using *bedtools*^68^ and 5’end of mapped coordinates extended 15 bp upstream and 22 bp downstream according to strand using *bedtools slop* (−l 15 -r 22 -s) parameters: to account for sterics during Tn5 transposition^77^. Replicates of extended coordinate BED files were concatenated and then converted back to BAM with *bedtools* and finally to bigWig using *deeptools bamCoverage*^70^ (parameters: --binSize 10 --normalizeUsing CPM --smoothLength 50 --extendReads 38). ATAC-seq peaks were called using *macs2 callpeak*^78^ (parameters= −f BAM, --keep-dup all --q 0.01).

### ChIP-seq processing

H3K27ac ChIP-seq reads were obtained from GEO (GSE118755)^16^ and mapped to the mm10 genome assembly with *bowtie2* with default parameters^75^. Mapped reads were filtered for mapping quality and PCR duplicates using *samtools view* and *markdup*^76^ (parameters: -q 30). Mapped reads from replicates were combined with *samtools merge*, extended according to sample and control average fragment estimates (“x”) from *macs2*^78^ and converted to bigWig signal tracks using *deeptools bamCompare* where control background signals (*e.g*., input) were subtracted from foreground (paramters: --operation subtract --binSize 50 -- scaleFactorsMethod None --normalizeUsing CPM --smoothLength 250 -- extendReads “x”).

### sc-RNA seq analysis

Counts data matrix were taken from (https://github.com/IStevant/XX-XY-mouse-gonad-scRNA-seq/tree/master/data/all_count.Robj)^73^. Cells of the clusters of interest were then extracted and divided randomly into two “pseudo-replicates”. Counts from these two pseudo-replicates were summed. Scaling and gene expression was then performed using the R package *DEseq2*^79^ and treated as bulk-RNA-seq with 2 replicates.

### RNA-seq bulk data processing

Reads were mapped using *STAR*^80^ (parameters: --outFilterMultimapNmax 1) to mm10 genome. Gene expression was quantified using *featureCounts* from the *subREAD* package^81^ with default parameters. Sample scaling and differential gene expression analysis was performed using the R package *DESeq2*^79^.

## Data availability

The Hi-C and bulk RNA-seq datasets generated in this study can be found in the Gene Expression Omnibus (GEO) with the accession code GSE217618.

The sc-RNA-seq counts matrix from supporting used populations was downloaded from https://github.com/IStevant/XX-XY-mouse-gonad-scRNA-seq/tree/master/data/all_count.Robj.

ChIP-seq and ATAC-seq raw fastq files were obtained from GEO (GSE118755).

## Code availability

All *METALoci* analysis was performed using in house developed Python 3 code available as a Jupyter notebook (https://github.com/3DGenomes). The rest of analysis were performed with previously published software packages or scripts, which are maintained and available in their respective repositories.

## ACKNOWLEDGMENTS

We thank the sequencing core, transgenic unit and animal facilities of the Max Delbrück Centre for Molecular for technical assistance. We thank C. Scholl, M. Altmann, G. Kussagk, S. Bomberg, S. Reissert-Oppermann and C. Westphal for their support with the transgenic work. We thank F. Martínez Real and members of the Lupiañez laboratory for their valuable input and comments on the manuscript. This research was supported by a grant from the Deutsche Forschungsgemeinschaft (International Research Training Group 2403) to I.M-G. and D.G.L. R.D.A. was supported by an EMBO Postdoctoral Fellowship (grant no. EMBO ALTF 537-2020). MAM-R acknowledges support by the Spanish Ministerio de Ciencia e Innovación (PID2020-115696RB-I00) and the National Human Genome Research Institute of the National Institutes of Health under Award Number RM1HG011016. B.C. and S.D. were supported by grants from the National Institute of Health (NIH R01-HD039963 and NIH R01-HD103064 to B.C.). The content is solely the responsibility of the authors and does not necessarily represent the official views of the National Institutes of Health.

## AUTHOR CONTRIBUTIONS

I.M-G., B.C., M.A.M-R., and D.G.L. conceived the study and designed the experiments. I.M-G. performed most experiments, with the support of A.H. S.D., and S.A.G-M. performed gonadal cell collections. J.A.R. and M.A.M-R conceptualized *METALoci* with the help of O.L. J.A.R. and M.A.M-R developed and applied *METALoci*. I.M-G and M.A.M-R performed most computational analyses, with the support of R.D.A. and S.L. I.M-G., M.A.M-R. and D.G.L. analyzed the data. J.J. and R.K. performed tetraploid aggregations. I.M-G., M.A.M-R. and D.G.L. wrote the manuscript with input from all authors.

