## Extended Data for "Sex-determining 3D regulatory hubs revealed by genome spatial auto-correlation analysis"

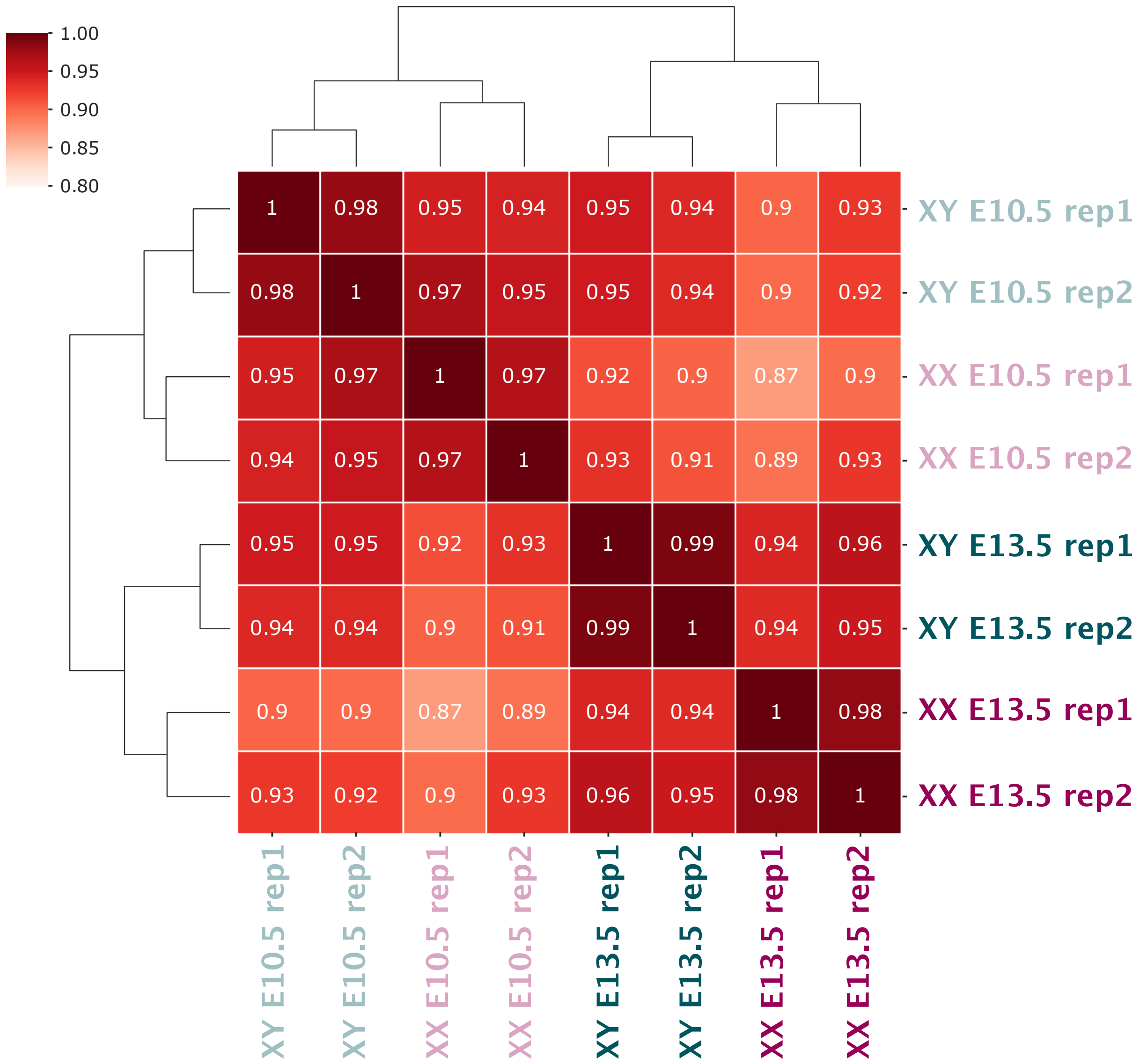


***Extended Data Fig. 1: Correlation of Hi-C samples based in A/B compartment signal.*** *Pearson correlation of the first eigenvector between samples and individual replicates. Note that replicates from each sample have increased correlation compared to other samples.*

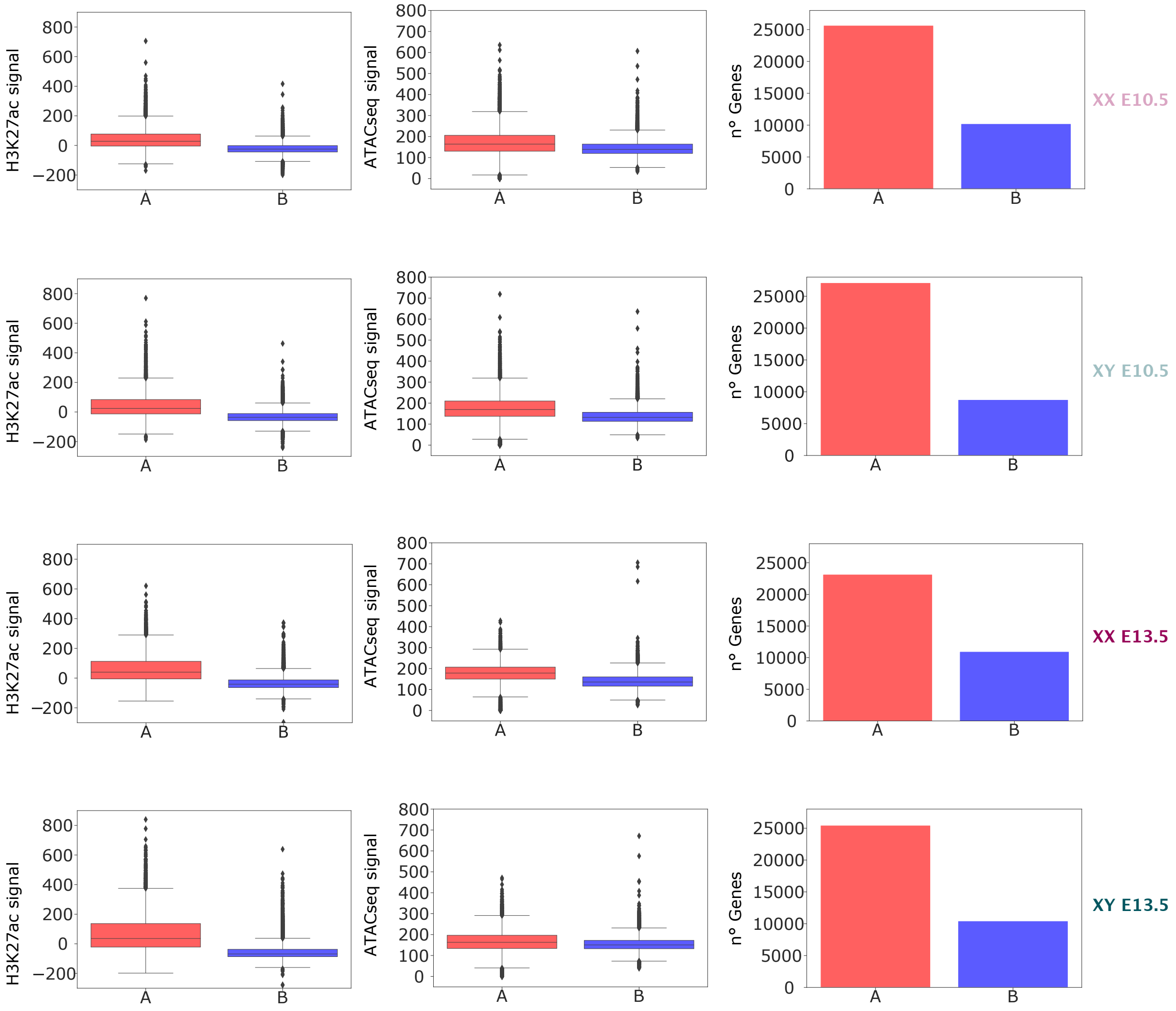


***Extended Data Fig. 2: Correlation of A/B compartments with active chromatin, open chromatin and with transcription.*** *H3K27ac ChIP-seq and ATAC-seq data obtained from[16].*

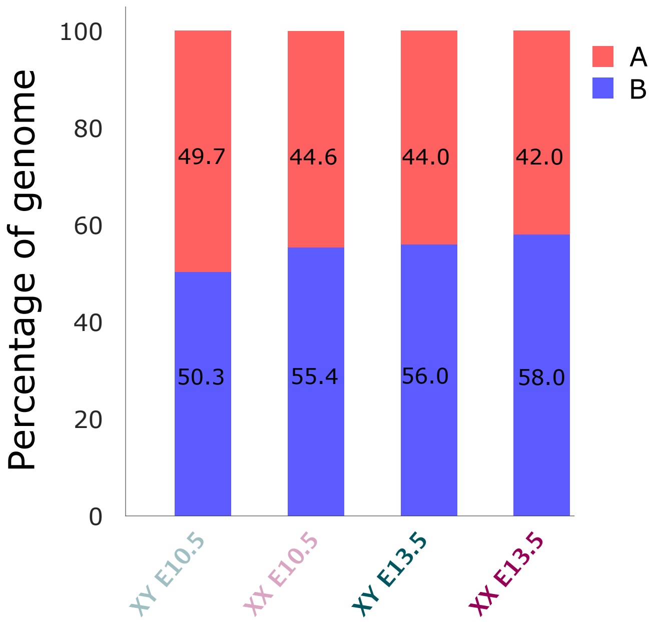


***Extended Data Fig. 3: Percentage of the genome located in A/B compartments.*** Stacked barplots representing which percentage of the genome corresponds to A compartment (red) or B compartment (blue) in each of the samples.

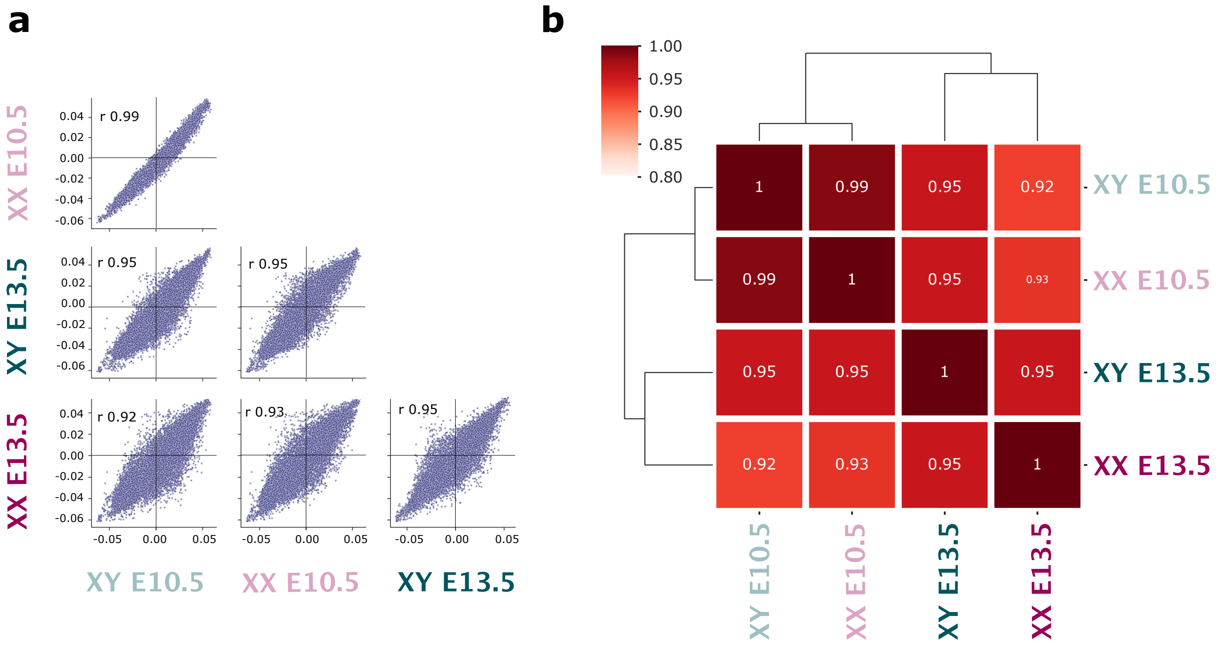


***Extended Data Fig. 4: Correlation of A/B compartments between different samples. a.*** *Scatterplot of pairwise correlation analysis of different samples.* ***b.*** *Heatmap of pairwise correlation analysis of different samples. For both graphs, note the increased correlation between XX E10.5 and XY E10.5, compared to other samples*

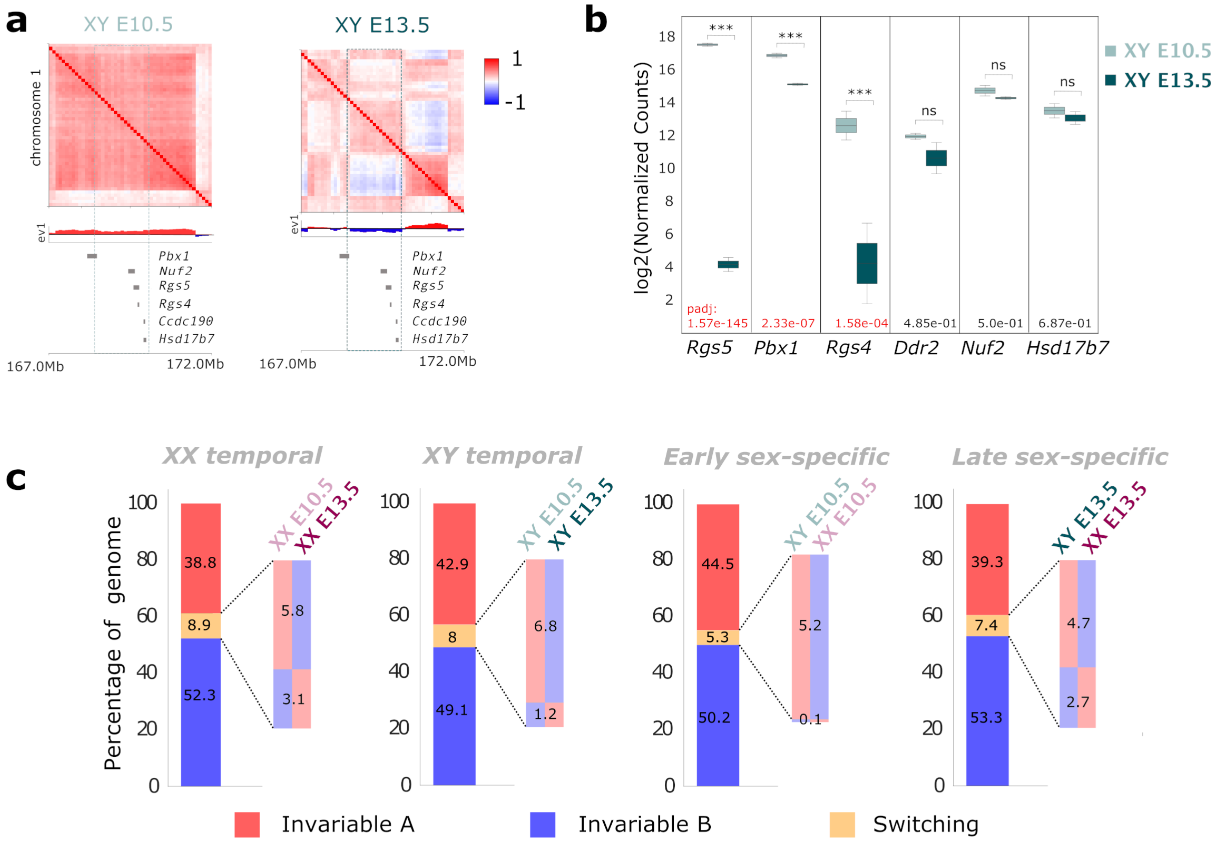


***Extended Data Fig. 5: A/B compartment dynamics during sex determination. a.*** *Example of a genomic regions that switch compartments during Sertoli cell differentiation.* ***b.*** *Expression levels of genes contained withing the A to B switched region in panel a. Note the decrease in expression for genes contained within the region.* ***c.*** *Percentage of genome that switches compartments or remain invariable.*

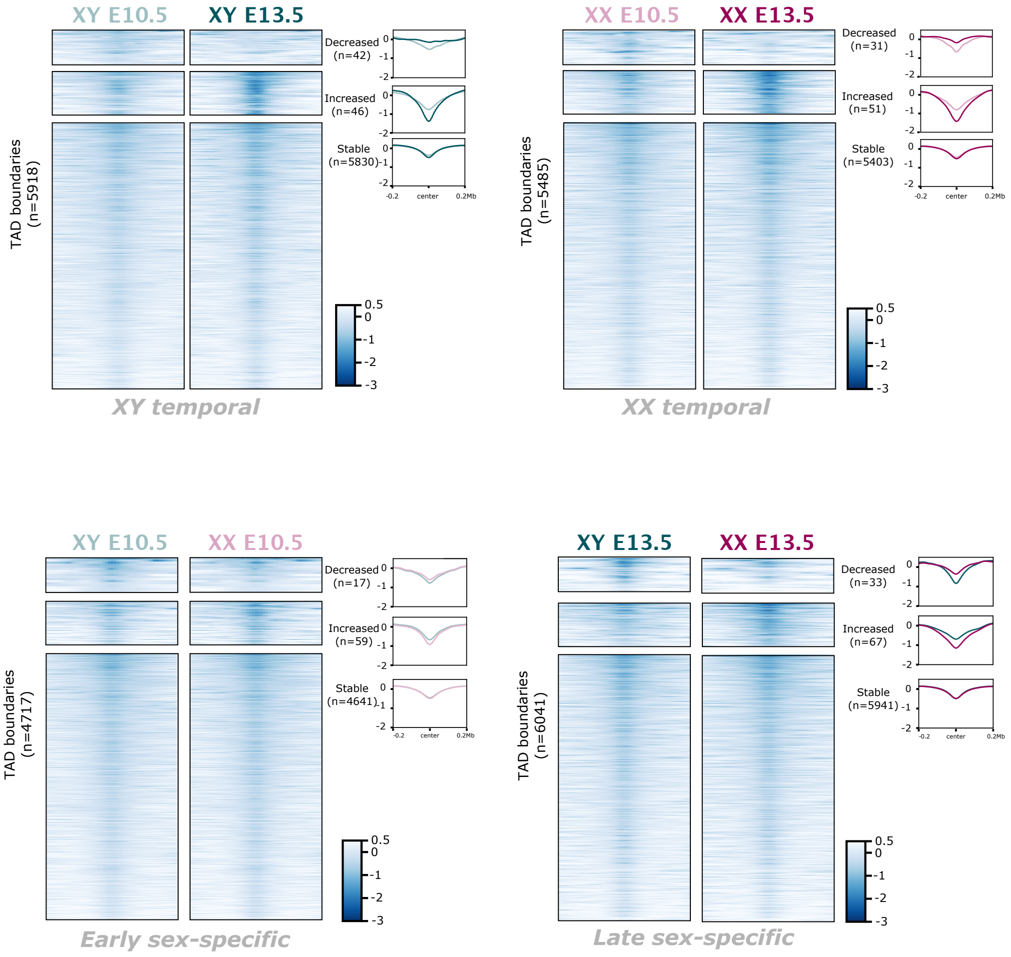


***Extended Data Fig. 6: Insulation heatmaps for TAD boundaries identified in pairwise comparisons.*** *Note that the "decreased" and "increased" heatmaps are enlarged with respect to the "stable" category, to facilitate visualization.*

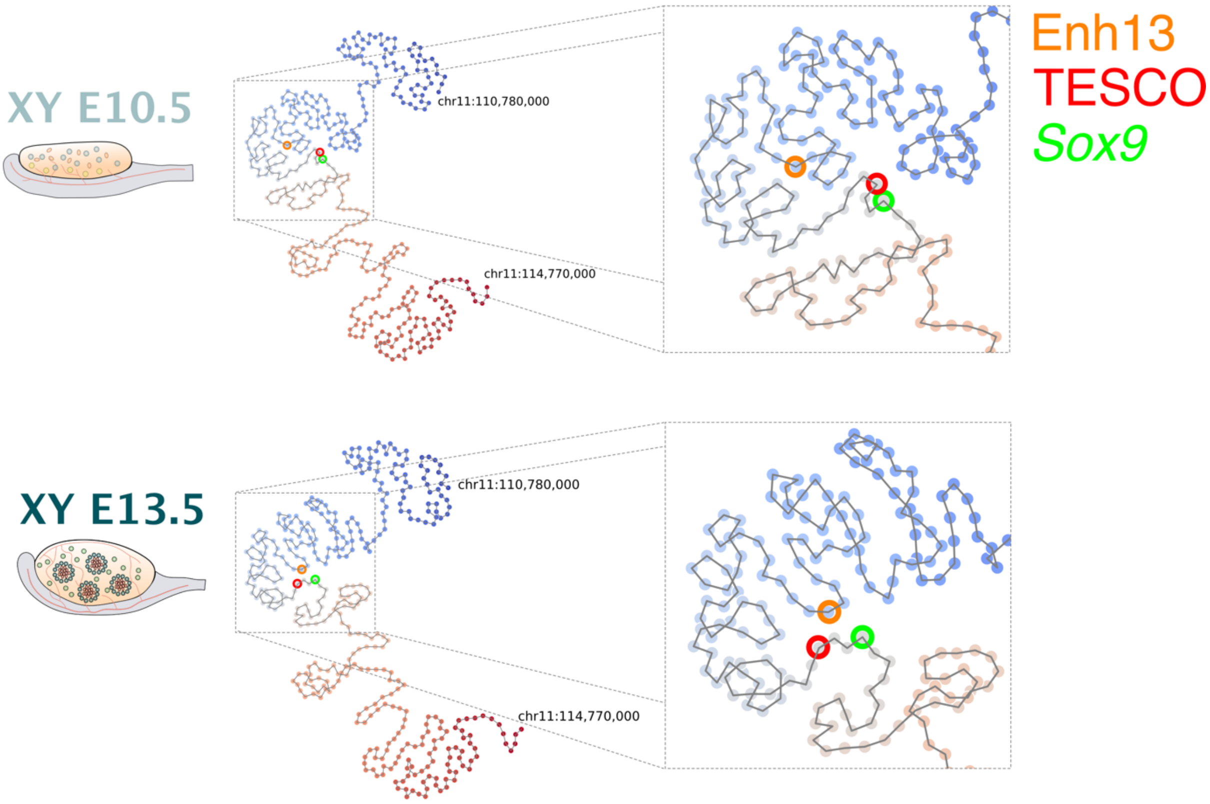


***Extended Data Fig. 7: Spatial dynamics of the* Sox9 *locus during Sertoli cell differentiation.*** *Left. Kamada-Kawai 2D layout of the* Sox9 *locus at XY E10.5 and E13.5 cells. Right panel. Zoom in of the squared region of the 2D layout. Note the increased proximity between Enh13 and the* Sox9 *promoter during Sertoli cell differentiation. The distance between the TESCO enhancer and the Sox9 promoter remain similar.*

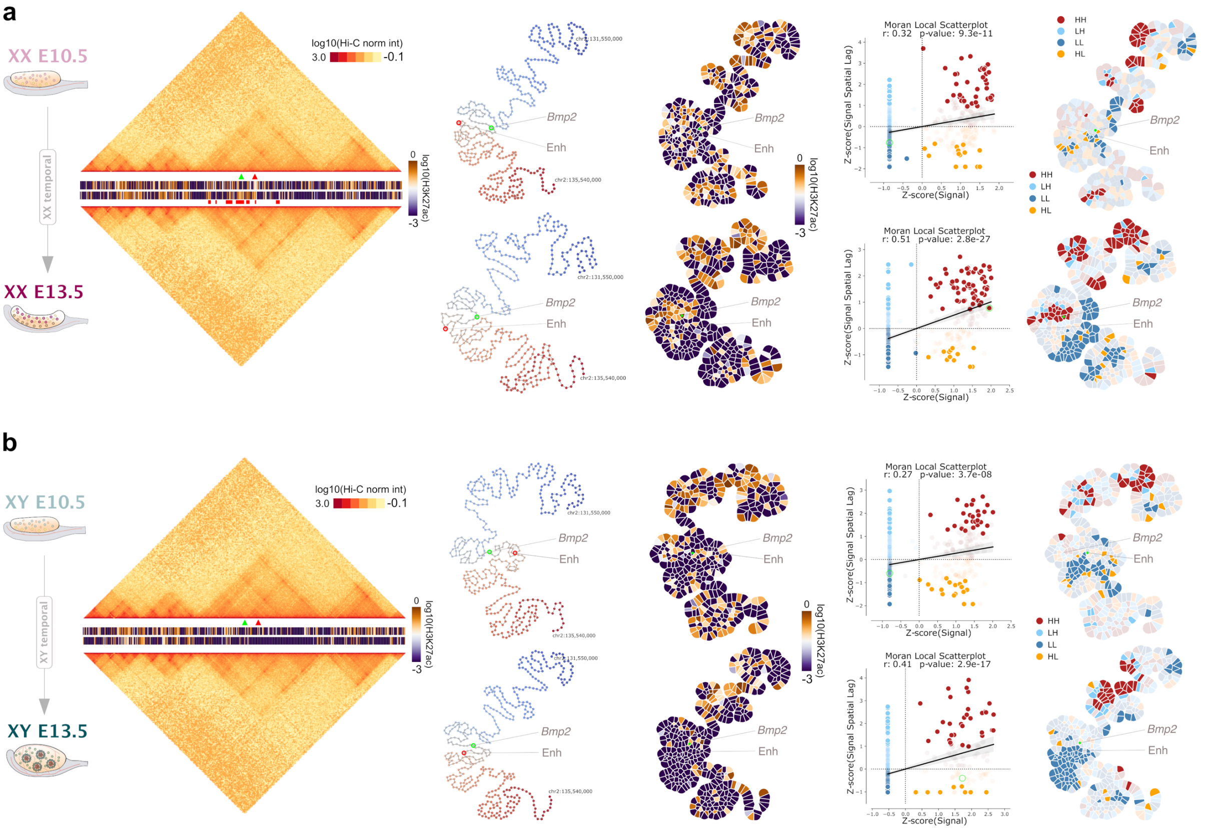


***Extended Data Fig. 8: METALoci analysis for the*** **Bmp2** ***locus during sex determination in both sexes. a*** *From left to right. First, Hi-C data and H3K27ac signal for* Bmp2 *locus centered at chr2:131,550,000-135,540,000 coordinates. Hi-C and H3K27ac ChIP-seq for granulosa cells development (XX E10.5 to XX E13.5). The position of the* Bmp2 *promoter (green arrowhead), as well as the recently identified enhancer*[16] *(red arrowhead), are highlighted. Squared red marks under H3K27ac track indicate the non-continous metaloci detected for* Bmp2 *locus at XX E13.5. Second, Kamada-Kawai 2D layout landmarking the position in the map for the* Bmp2 *promoter (green circle) as well as the annotated enhancer (red circle). Third, H3K27ac signal mapped into the graph layout and represented as a Gaudí plot. Fourth, LMI scatter plot. And fifth, Gaudí plot highlighting the LMI quadrants with solid color indicating statistical significance (p<0.05).* ***d.*** *Same as* ***a*** *for Sertoli development (XY E10.5 to XY E13.5 cells).*

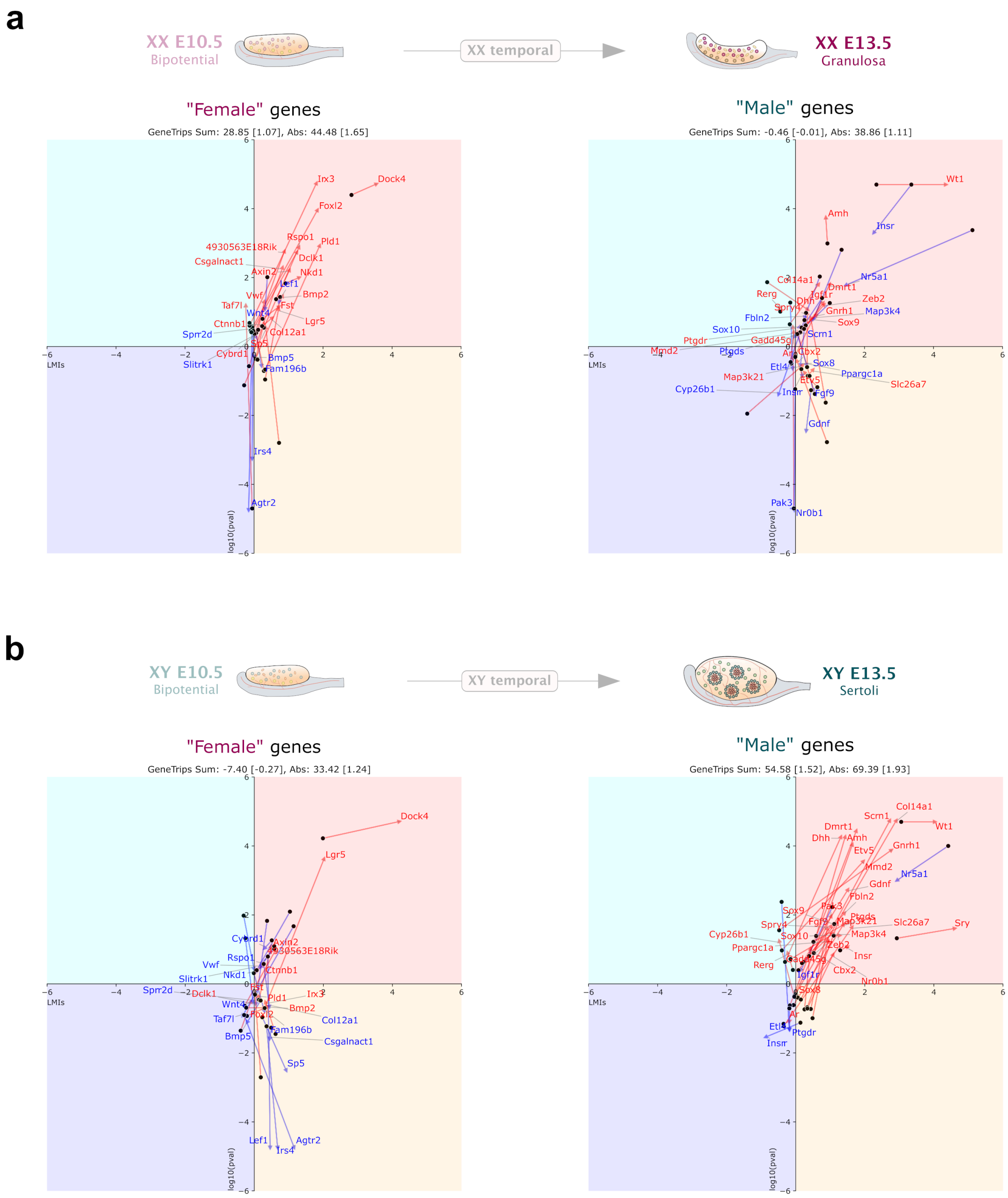


***Extended Data Fig. 9: Gene trips for male- and female-specific genes during sex determination in both sexes. a.*** *LMI transition (or "gene trips") for genes acquiring female and male-specific expression during sex differentiation for the differentiation of granulosa cells (XX E10.5 to XX E13.5). The plots show the four LMI quadrans (that is, HH, LH, LL, and HL in the LMI p-value and LM's I axis). Arrows indicate individual trip for each gene, which names are indicated at the arrowhead. Red arrows indicate tendency to trip towards H3K27ac environment while blue arrows indicate otherwise. Gene trip sum as well as the sum of absolute gene trips are shown at the top of the graph together with mean values per gene shown between brackets.* ***b****. Same as* ***a*** *for the differentiation of Sertoli cells (XY E10.5 to XY E13.5).*

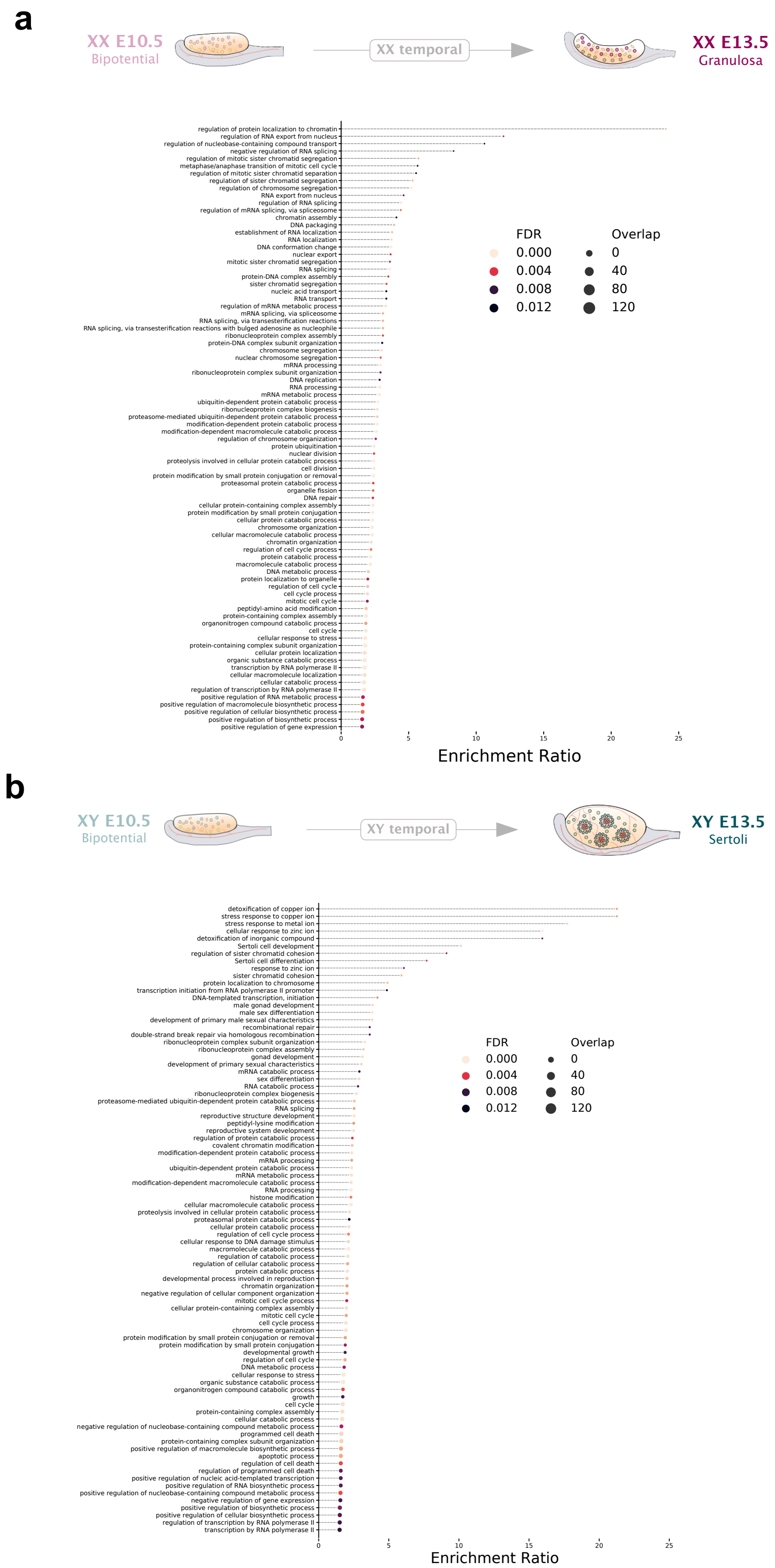


***Extended Data Fig. 10: Full list of GO enrichment terms during sex determination in both sexes. a.*** *GO Biological Process enrichment analysis for genes that transition from LL or HL to HH during granulosa cell differentiation (XX E10.5 to XX E13.5).* ***b.*** *Same as* ***a*** *for Sertoli cell differentiation (XY E10.5 to XY E13.5).*

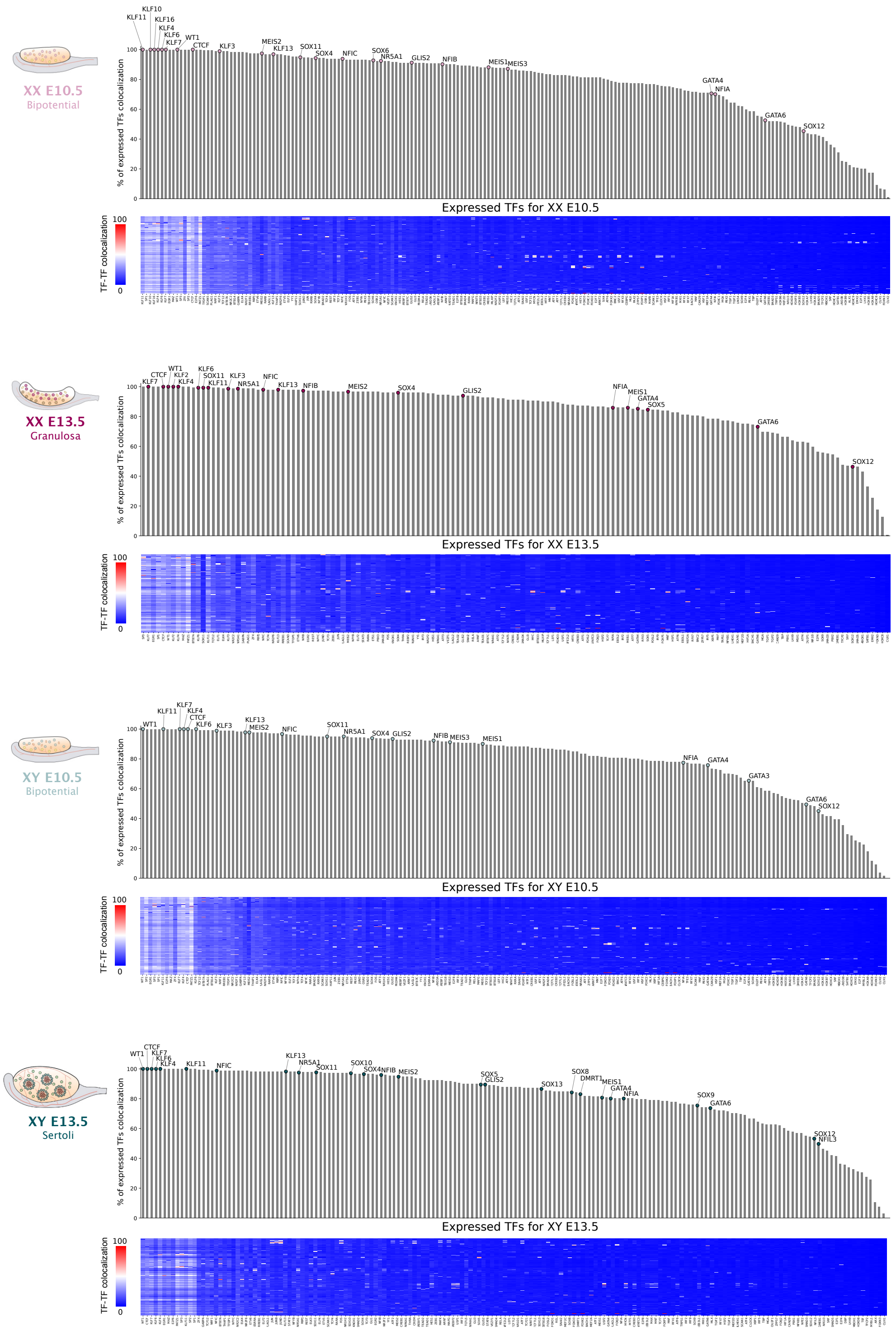


***Extended Data Fig. 11: STRIPE analysis for colocalization of TFs.*** *For each cell type, the bar graph shows all expressed TFs in the cell type sorted by the percentage of TFs that colocalize with them. A TF is considered to colocalize if both are found together in the genome at least 20 instances. Relevant TFs for sex-determination are highlighted. A total of 198, 153, 186, and 176 TFs are expressed in XX E10.5, XX E13.5, XY E10.5, and XY E13.5, respectively*[73].

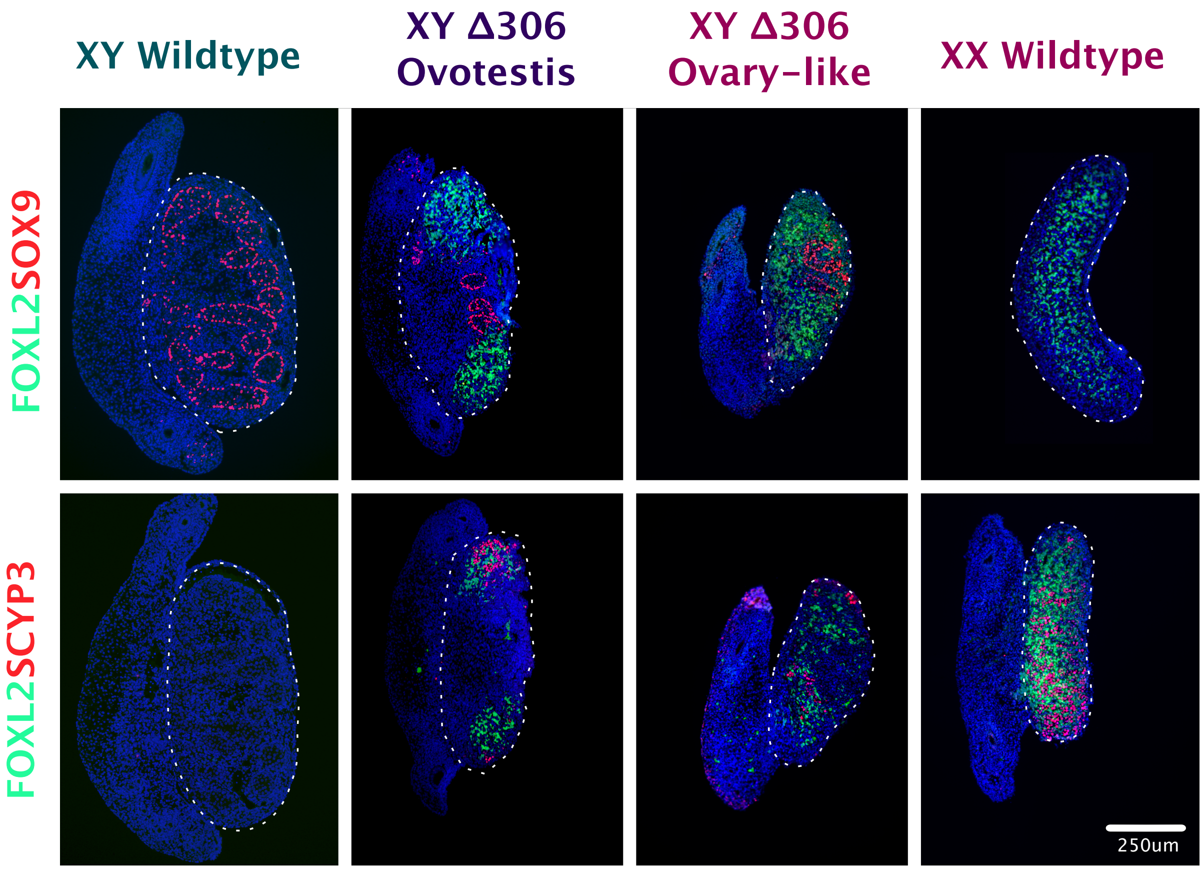


***Extended Data Fig. 12: Immunofluorescence analysis for sex-specific and meiotic markers .*** *Upper panel shows immunofluorescence for the testis marker SOX9 (red) and the ovarian marker FOXL2 (green). Note the presence of both markers in the mutant gonad. Lower panel show immunofluorescence of female marker FOXL2 (green) and meiosis marker SYCP3 (red), note of the presence of both markers in XY Δ306 and XX wild type gonads (middle and right) and the absence of them in the XY wild type gonad (left). Gonads are delineated by a discontinuous line.*

***Extended Data File 1: Coordinates for HH metaloci in each sample.***

*The EDFile1_H3K27ac_metaloci_per_gene.zip file contains four files named:*

- XX10.5_H3K27ac_HH_metaloci_per_gene.bed
- XX13.5_H3K27ac_HH_metaloci_per_gene.bed
- XY10.5_H3K27ac_HH_metaloci_per_gene.bed
- XY13.5_H3K27ac_HH_metaloci_per_gene.bed

*Each BED file contains the following columns tab separated:*

| | |
|---|---|
| chr | Chromosome |
| start | Start coordinates |
| end | End coordinates |
| MetaLociBinNumber | Number of the bin in the METALoci layout |
| GeneSymbol | Gene symbol |

*This ZIP file can be anonymously accessed in ZENODO (10.5281/zenodo.7334816) at this URL: https://doi.org/10.5281/zenodo.7334816*

***Extended Data File 2: Coordinates for all TFs assessed by TOBIAS to bind an ATAC-seq accessible peak within a metaloci in each sample.***

*The EDFile2_TF_ATAC_ML_Sites.zip file contains four files named:*

- XX10.5_TF_ATAC_ML_Sites.tsv
- XX13.5_TF_ATAC_ML_Sites.tsv
- XY10.5_TF_ATAC_ML_Sites.tsv
- XY13.5_TF_ATAC_ML_Sites.tsv

*Each TSV file contains the following columns tab separated:*

| | |
|---|---|
| chr | Motif site chromosome |
| start | Motif site start coordinates |
| end | Motif site end coordinates |
| tfscore | TOBIAS transcription factor score |
| strand | Motif site strand |
| pchr | ATAC site chromosome |
| pstart | ATAC site start coordinates |
| pend | ATAC site end coordinates |
| samplescore | TOBIAS sample score |
| bchr | METALoci chromosome |
| bstart | METALoci start coordinates |
| bend | METALoci end coordinates |
| bin | METALoci bin |
| gene | Gene symbol |
| tf | Transcription Factor |
| motif | Motif id |

*This ZIP file can be anonymously accessed in ZENODO (10.5281/zenodo.7334816) at this URL: [https://doi.org/10.5281/zenodo.7334816](https://doi.org/10.5281/zenodo.7334816)*
